# DipM controls multiple autolysins and mediates two regulatory feedback loops promoting cell constriction in *C. crescentus*

**DOI:** 10.1101/2022.09.10.507391

**Authors:** Adrian Izquierdo-Martinez, Vega Miguel-Ruano, Rogelio Hernández-Tamayo, Jacob Biboy, María T. Batuecas, Maria Billini, Timo Glatter, Waldemar Vollmer, Peter L. Graumann, Juan A. Hermoso, Martin Thanbichler

**Author notes:** Address correspondence to Martin Thanbichler. Bacterial Cell Biology, Instituto de Tecnologia Química e Biológica António Xavier, Universidade Nova de Lisboa, Oeiras, Portugal (Adrian Izquierdo-Martinez).

## Abstract

Proteins containing a catalytically inactive LytM-type endopeptidase domain have emerged as important regulators of cell wall-degrading enzymes in bacteria. Although these so-called LytM factors are wide-spread among species, the range of functions they fulfill and their precise modes of action are still incompletely understood. In this work, we study the LytM factor DipM, a protein required for proper cell division in the model species *C. crescentus*. We show that the LytM domain of DipM interacts directly with multiple autolysins, including the lytic transglycosylases SdpA and SdpB, the amidase AmiC and the putative carboxypeptidase CrbA, and stimulates the activities of SdpA and AmiC. The crystal structure of the LytM domain of DipM reveals conserved features, including a distinctive groove. Modeling studies suggest that this groove could represent the docking site of AmiC. The architecture of the binding interface in the DipM-AmiC complex is very similar to that observed for the LytM domain of EnvC in complex with its autoinhibitory restraining arm, suggesting a conserved role of the groove in the interaction of LytM factors with their (auto-)regulatory targets. In line with this hypothesis, a mutation in the groove abolishes DipM function. Interestingly, single-molecule tracking studies reveal that the recruitment of DipM and its regulatory targets SdpA and SdpB to the division site is mutually interdependent, with DipM establishing a self-reinforcing cycle that gradually increases lytic transglycosylase activity at the cell center as division progresses. At the same time, the DipM-dependent activation of AmiC leads to the production of denuded peptidoglycan, generating a spatial cue that attracts FtsN to the division site and thus, in turn, again promotes the recruitment of DipM. Collectively, these findings show that DipM is a central regulator that acts at the intersection of different peptidoglycan remodeling pathways and coordinates the activities of various classes of autolysins to promote cell constriction and daughter cell separation.

## Introduction

In the course of evolution, cells have developed multiple strategies to reinforce their envelope in order to resist the internal osmotic pressure. Most bacterial species synthesize a semi-rigid cell wall surrounding the cytoplasmic membrane that bears part of the tension force generated and, in turn, gives shape to the cells. The central component of the bacterial cell wall is peptidoglycan (PG) (Schleifer and Kandler, 1972; Weidel et al., 1960), a heteropolymer composed of glycan strands of alternating N-acetylglucosamine (NAG) and N-acetylmuramic acid (NAM) units that are covalently crosslinked by short peptide bridges (Vollmer et al., 2008). The PG meshwork constitutes a single large macromolecule, the so-called sacculus, which needs to be constantly remodeled to enable cell growth, morphogenesis and cell division. This process requires the cleavage of bonds within the sacculus by lytic enzymes, also known as autolysins, and the subsequent insertion of new cell wall material by PG synthases. The activities of these two antagonistic groups of proteins need to be closely coordinated to prevent the emergence of weak spots in the PG layer that result in cell lysis (Höltje, 1998).

Autolysins are a heterogeneous group of enzymes that are classified according to the bond they break in the PG molecule. Glycosidases and lytic transglycosylases (LTs) cleave the bonds between the sugar units of the glycan stands (Dik et al., 2017). Notably, the reaction mediated by LTs produces 1,6-anhydro-NAM, which in some species acts as a signaling molecule indicating β-lactam antibiotic stress (Jacobs et al., 1994). N-acetylmuramyl-L-alanine amidases (PG amidases) hydrolyze the bond between the L-alanine residue of the peptide and the lactyl moieties of NAM, generating so-called naked glycan strands. They have been found to be required for daughter cell separation in various members of the gammaproteobacteria and firmicutes (Heidrich et al., 2001; Möll et al., 2014; Reinscheid et al., 2001; Yakhnina et al., 2015). Finally, endopeptidases break various bonds in the peptide moieties, promoting PG incorporation and remodeling (Chodisetti and Reddy, 2019; Firczuk and Bochtler, 2007; Hashimoto et al., 2012; Singh et al., 2012). In general, individual autolysins are rarely essential. However, in many bacteria, the combined inactivation of multiple autolysins can cause strong morphological and/or synthetic lethal phenotypes (Dörr et al., 2013; Hashimoto et al., 2012; Heidrich et al., 2002; Martinelli and Pavelka, 2016; Zielińska et al., 2017).

The coordination of lytic and synthetic enzymes is thought to be achieved by their assembly into multi-protein complexes (Höltje, 1998; Pazos et al., 2017; Typas et al., 2011). One of these complexes is the divisome, which carries out cell division in most bacteria (Lutkenhaus, 2009; Typas et al., 2011). Its assembly typically initiates with the polymerization of the tubulin homolog FtsZ into a dynamic ring-like structure at the future division site (Mukherjee and Lutkenhaus, 1994; Ortiz et al., 2016). This so-called Z-ring then recruits, directly or indirectly, all other divisome components, including PG synthases, autolysins and regulatory proteins (Du and Lutkenhaus, 2017; Goley et al., 2011). The coordinated activity of these factors gradually remodels the PG layer at the division site, constricting and ultimately splitting the sacculus to enable the release of the nascent daughter cells.

One of the most conserved mechanisms to regulate autolysin activity relies on the divisome components FtsE and FtsX, which form an ABC transporter-like complex in the cytoplasmic membrane (Schmidt et al., 2004). It is thought that the ATPase activity of FtsE induces conformational changes in the transmembrane protein FtsX, which then conveys this signal to specific autolysins, thereby controlling their activity state (Arends et al., 2009; Bartual et al., 2014; Lim et al., 2019; Mavrici et al., 2014; Meisner et al., 2013; Rued et al., 2019; Sham et al., 2011). In the case of gammaproteobacteria, this regulatory cascade involves endopeptidase homologs with a catalytically inactive LytM domain (termed *LytM factors*), which act as adaptors linking FtsX to PG amidases (Möll et al., 2014; Uehara et al., 2009; Uehara et al., 2010; Yakhnina et al., 2015; Yang et al., 2012). Importantly, there are also several examples of LytM factors that control autolysin activity in an FtsEX-independent manner (Möll et al., 2014; Tsang et al., 2017; Uehara et al., 2010; Yakhnina et al., 2015; Yang et al., 2018). Their regulatory targets and modes of action are highly variable among species, making it difficult to draw firm conclusions about the roles of LytM factors in previously uncharacterized systems.

*Caulobacter crescentus* is a crescent-shaped Gram-negative bacterium that serves as one of the primary model organisms to study cell cycle regulation, cell differentiation and morphogenesis in the alphaproteobacteria (Curtis and Brun, 2010; Henrici and Johnson, 1935; Poindexter, 1981). It is characterized by a dimorphic life cycle in which a newborn motile swarmer differentiates into a sessile and division-competent stalked cell. The stalked cell then elongates and divides asymmetrically, giving rise to a swarmer and a stalked sibling (Curtis and Brun, 2010). During the swarmer cell stage, the cell body grows in length by disperse incorporation of new PG in the lateral cell walls. This process is mediated by the so-called elongasome, a multi-protein complex controlled by the actin homolog MreB (Dye et al., 2005; Gitai et al., 2004). After the initiation of divisome assembly, the *C. crescentus* cell switches to a medial mode of growth and, in addition, initiates stalk biosynthesis at its old pole (Aaron et al., 2007; Billini et al., 2019; Kühn et al., 2010; Wagner et al., 2005). Finally, activation of the divisome leads to progressive constriction of the mid-cell region, which culminates with the separation of the two nascent daughter cells.

Previous studies have comprehensively analyzed the autolytic machinery of *C. crescentus* and the contributions of its components to cell morphogenesis in this species. A key regulatory factor is the LytM factor DipM (Goley et al., 2010; Möll et al., 2010; Poggio et al., 2010), a soluble periplasmic protein that carries two tandems of PG-binding LysM domains, and a C-terminal catalytically inactive LytM domain. The LysM domains are required for the accumulation of DipM at the division site, which initiates the early stages of divisome assembly (Goley et al., 2010; Goley et al., 2011; Möll et al., 2010; Poggio et al., 2010). Deletion of *dipM* causes a pronounced filamentation phenotype, accompanied by membrane blebbing, although the published data do not completely agree on the degree of severity (Goley et al., 2010; Möll et al., 2010; Poggio et al., 2010). Interestingly, the LytM domain alone appears to be sufficient to complement the Δ*dipM* phenotype, indicating that this domain most probably carries out the regulatory activity that DipM may have (Goley et al., 2010; Möll et al., 2010; Poggio et al., 2010). Previous work has shown that DipM is required, directly or indirectly, to recruit the putative soluble lytic transglycosylases (SLTs) SdpA and SdpB to the division site (Zielińska et al., 2017). Moreover, *in vitro* studies revealed that it is able to stimulate the enzymatic activity of AmiC, the essential putative PG amidase of *C. crescentus* (Meier et al., 2017; Zielińska et al., 2017). Notably, besides DipM, *C. crescentus* contains another soluble LytM factor, named LdpF, which features two N-terminal coiled-coil domains and has a global architecture similar to that of *E. coli* EnvC (Zielińska et al., 2017). LdpF is required for the mid-cell recruitment of AmiC, but *in vitro* evidence suggests that, in contrast to DipM, it is unable to stimulate the catalytic activity of AmiC (Meier et al., 2017; Zielińska et al., 2017). Unlike the two LytM factors, the catalytically active LytM endopeptidases (LdpA-E) of *C. crescentus* are largely redundant and only contribute to general cell fitness (Zielińska et al., 2017). Notably, DipM and SdpA, together with the D,D-carboxypeptidase CrbA, are also recruited to the stalked pole, where they form part of a distinct PG biosynthetic complex mediating stalk elongation (Billini et al., 2019). In conclusion, genetic, protein localization and *in vitro* studies have provided solid, but so far only fragmentary information about the regulatory roles of DipM and LdpF, which is insufficient to comprehensively understand the physiological roles of LytM factors in *C. crescentus*.

In the present study, we characterized the interactome of the *C. crescentus* LytM factors DipM and LdpF using various *in vivo* and *in vitro* approaches. Our results indicate that DipM interacts with four different autolysins (SdpA, SdpB, AmiC, CrbA) and the divisome component FtsN *in vivo*. We verified these interactions *in vitro* and revealed that the five interactors compete for DipM and do not produce ternary or higher-order complexes. Moreover, using PG degradation assays, we obtained direct evidence for a stimulatory effect of DipM on the lytic activities of SdpA and AmiC. To improve our mechanistic understanding of DipM, we solved the crystal structure of the LytM domain of DipM and thus identified DipM-specific features as well as potential interaction interfaces that may serve as docking sites for its different interactors. Finally, single-molecule tracking studies reveal the existence of a self-reinforcing cycle, in which the DipM-dependent mid-cell recruitment of SdpA and SdpB in turn increases the residence time of DipM in this location, leading to a progressive increase in autolytic activity as cell constriction proceeds. Collectively, our results demonstrate that *C. crescentus* DipM coordinates a complex autolysin network whose topology greatly differs from that of previously investigated autolysin systems.

## Results

### DipM mediates a novel interaction network

Previous studies have revealed that DipM mediates the mid-cell localization of the SLTs SdpA and SdpB (Zielińska et al., 2017) and stimulates the activity of the PG amidase AmiC (Meier et al., 2017; Zielińska et al., 2017). However, the phenotype of a Δ*dipM* mutant, characterized by severe filamentation, membrane blebbing and ectopic pole formation, is more severe and pleiotropic than that of a Δ*sdpA* Δ*sdpB* double mutant, which only shows mild filamentation (Zielińska et al., 2017). Moreover, the Δ*dipM* phenotype also clearly differs from the one observed upon depletion of AmiC, i.e. cell chaining (Meier et al., 2017; Zielińska et al., 2017). Therefore, we concluded that DipM must be involved in additional pathways.

In order to fully elucidate the function of DipM, we determined its interactome in live cells using co-immunoprecipitation (Co-IP) analysis. To this end, we constructed a strain that produced a fully functional DipM variant carrying a C-terminal FLAG affinity peptide (DipM-FLAG) in place of the wild-type protein. After crosslinking with formaldehyde, protein complexes were captured with anti-FLAG affinity beads and subjected to mass spectrometry. Several proteins were specifically enriched in this analysis (**Figure 1A**), including PG-degrading enzymes (SdpA, SdpB, AmiC, CrbA), the cell division protein FtsN, the PG synthase PbpX (Strobel et al., 2014) and various so-far uncharacterized proteins. For the present study, we decided to focus on SdpA, SdpB, AmiC, CrbA and FtsN, which, like DipM, have been shown to be involved in cell division and/or stalk elongation (Billini et al., 2019; Möll and Thanbichler, 2009; Zielińska et al., 2017). To confirm the interactions *in vivo*, we performed reciprocal Co-IP analyses using the identified proteins as baits. We observed a specific enrichment of DipM by FLAG-tagged versions of SdpA (**Figure 1B**), FtsN (**Figure 1C**) and CrbA (**Figure 1D**), corroborating their interaction. In a complementary analysis, a similar set of experiments was conducted on the second LytM factor of *C. crescentus*, LdpF (Meier et al., 2017; Zielińska et al., 2017). The results obtained indicate a mutual interaction of AmiC and LdpF, and they provide evidence for the interaction of both proteins with the FtsEX complex (**Figure 1E-F**). Collectively, these findings suggest the existence of two different regulatory hubs (Fig. 1G). The first one, DipM, interacts with the lytic enzymes SdpA, SdpB, AmiC and CrbA as well as the divisome component FtsN, a bitopic membrane protein serving as a central activator of PG remodeling enzymes at the division site (Weiss, 2015). The second hub, LdpF, by contrast, likely connects AmiC with the FtsEX complex, further supporting the notion that it represents a functional homolog of the gammaproteobacterial EnvC proteins.

**Figure 1.**
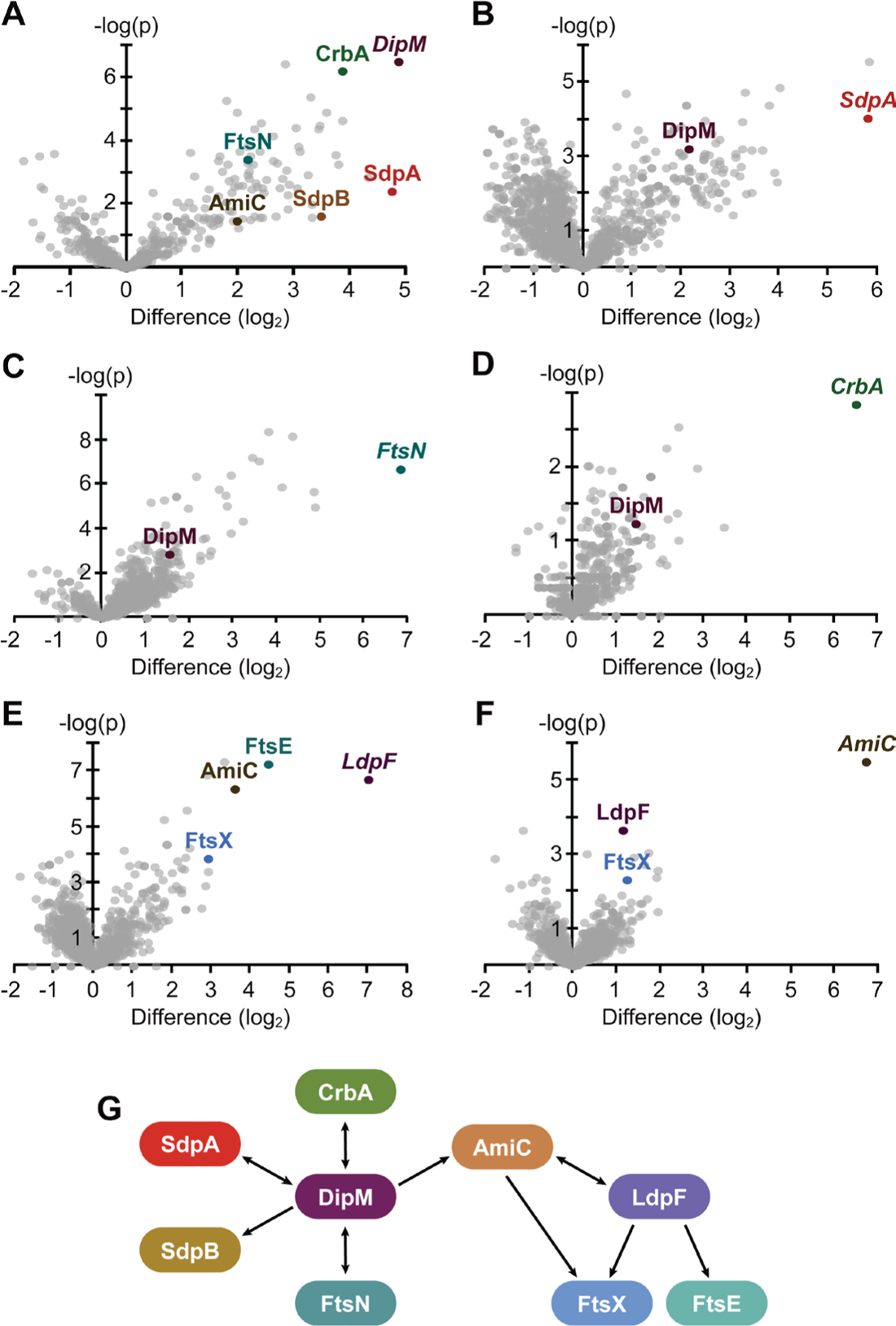
Interactome of DipM and LdpF. **(A-F)** Volcano plots showing the interactors of **(A)** DipM-FLAG (four replicates), **(B)** SdpA-FLAG (three replicates), **(C)** FtsN-GFP (four replicates), **(D)** CrbA-FLAG (two replicates), **(E)** LdpF-FLAG (four replicates) and **(F)** AmiC-FLAG (four replicates) as identified by Co-IP analysis. The data points show the log2 of the average differences in the peptide counts for each hit compared to the control (x-axis) plotted against the –log10 of the *p* values of these peptide counts (y-axis). Dots in color represent hits relevant for the purpose of this study, with the names of the corresponding proteins given in the same color next to them. Italic font is used for the bait proteins used in the respective experiment. **(G)** Schematic representation of the relevant hits and their interactions as identified by Co-IP analysis. Arrows point from the bait proteins to the corresponding enriched prey proteins.

To determine whether the interactions observed by co-immunoprecipitation analysis were direct, we purified the proteins or soluble fragments of them and analyzed their interaction by bio-layer interferometry (BLI). To this end, the presumed DipM targets were immobilized on BLI sensors and probed with DipM as an analyte. The results obtained showed that DipM associates with SdpA, SdpB, AmiC, the SPOR domain of CrbA and the periplasmic region of FtsN (**Figure 2B-E** and **Figure 3**). In all cases, the complexes were highly dynamic, with fast association and dissociation kinetics and equilibrium dissociation constants (*K_D_*) in the low micromolar range (3-50 μM).

**Figure 2.**
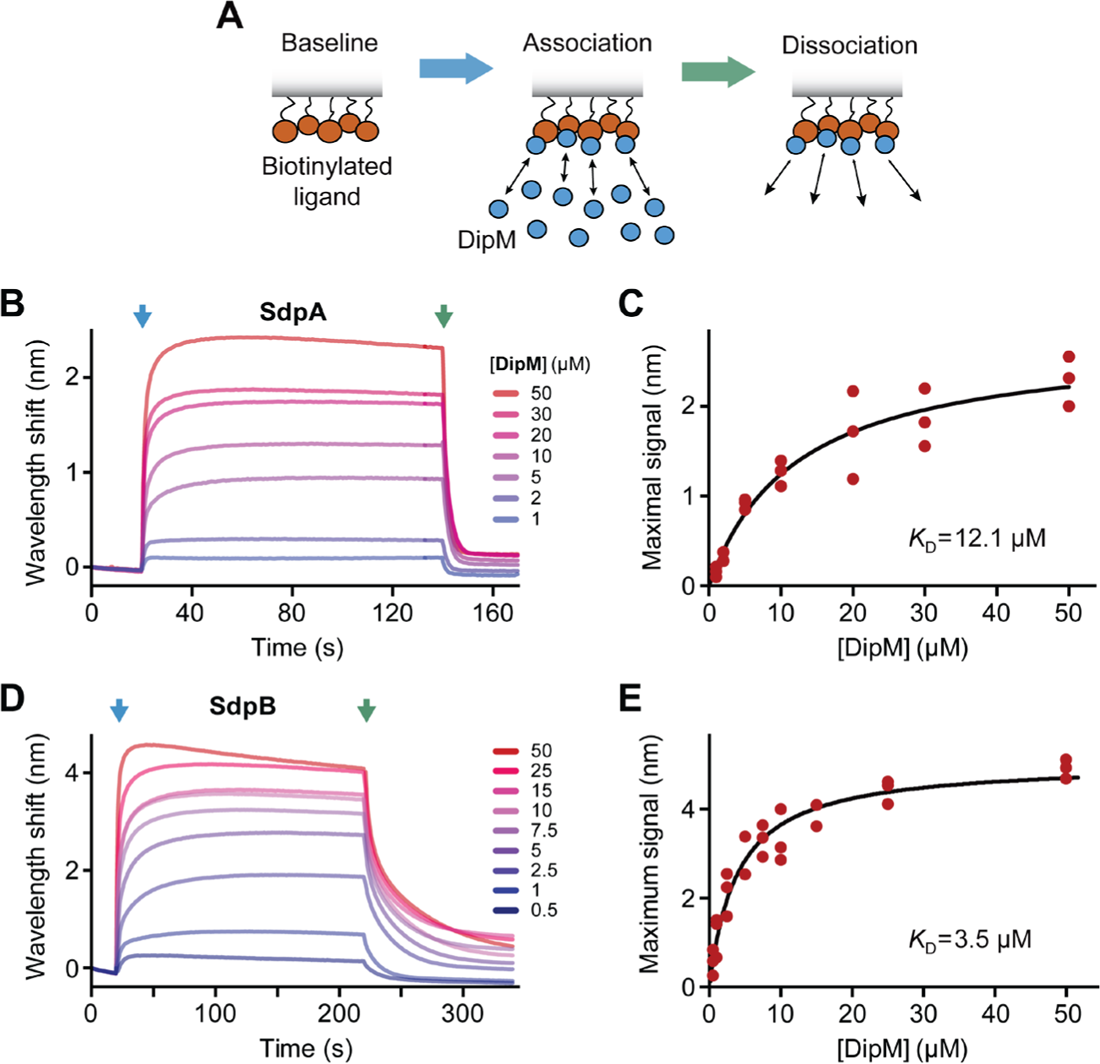
DipM binds SdpA and SdpB *in vitro*. **(A)** Schematic showing the different steps of a bio-layer interferometry (BLI) experiment. After immobilization of the biotinylated ligand, the sensor is washed with buffer to establish a stable baseline. To start the association reaction, the sensor is dipped into a solution of the analyte prepared in the same buffer, which will progressively bind to the immobilized ligand until equilibrium is reached. Finally, the sensor is dipped into analyte-free buffer to monitor the dissociation reaction. **(B)** Representative BLI experiment showing the interaction of DipM (at the indicated concentrations) with immobilized SdpA. The blue arrow marks the beginning of the association phase, the green arrow the beginning of the dissociation phase. **(C)** Graph showing the wavelength shifts after equilibration of the binding reactions plotted against the corresponding DipM concentrations in BLI experiments using sensors functionalized with SdpA (n=3). The blue line represents the best fit to a one-site binding model. **(D, E)** Analogous to panels B and C, respectively, but using sensors functionalized with biotinylated SdpB.

**Figure 3.**
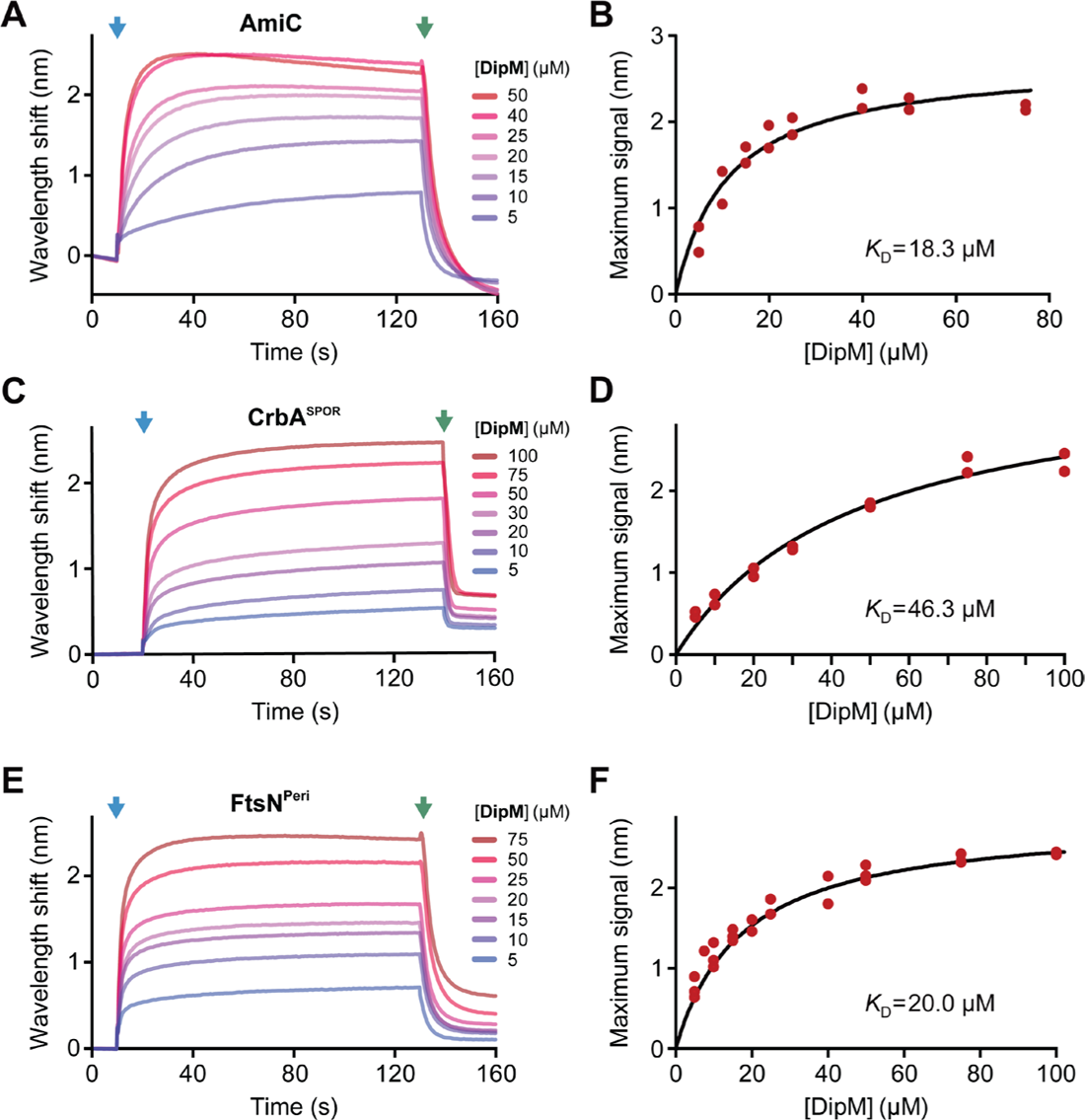
DipM binds FtsN, AmiC and CrbA *in vitro*. **(A)** Representative BLI experiment showing the interaction of DipM (at the indicated concentrations) with immobilized AmiC. The blue arrow marks the beginning of the association phase, the green arrow the beginning of the dissociation phase. **(B)** Graph showing the wavelength shifts after equilibration of the binding reactions plotted against the corresponding DipM concentrations in BLI experiments using sensors functionalized with AmiC (n=3). The blue line represents the best fit to a one-site binding model. **(C,D)** Analogous to panels A and B, respectively, but using sensors functionalized with the biotinylated SPOR domain of CrbA (CrbA^SPOR^). **(E,F)** Analogous to panels A and B, respectively, but using sensors functionalized with a biotinylated fragment comprising the periplasmic region of FtsN (FtsN^Peri^).

We also considered the possibility that DipM acts as an interaction platform that can simultaneously bind multiple different target proteins. Therefore, we tested whether the preincubation of DipM with a second factor before its addition to the BLI sensors resulted in an increase in the binding signal, indicating the formation of a ternary complex, or rather in a decrease of the signal, which would suggest a competition of the DipM target proteins for the same binding site (**Figure S1A**). It was not possible to analyze all combinations due to non-specific binding of some of the proteins to the sensor surface. However, for all protein pairs analyzed, the signals remained largely unchanged or decreased compared to the reactions with DipM alone (**Figure S1B-E**). These findings indicate that the target proteins compete for binding to DipM or, conversely, that DipM does not mediate the formation of a multi-protein complex under the conditions tested.

### DipM enhances the activities of SdpA and AmiC *in vitro*

Having confirmed a direct interaction of DipM with various autolysins, we next investigated the possibility that DipM could have a regulatory effect on the enzymatic activity of its target proteins. To clarify this point, we performed PG digestion assays with purified enzymes in the absence or presence of DipM (**Figure 4A**). When incubated with purified sacculi, SdpA alone showed basal lytic activity against non-crosslinked and, to a small extent, also crosslinked PG, producing muropeptides that contained 1,6-anhydro-MurNAc moieties (**Figure 4B**). These data confirm the bioinformatic prediction that SdpA is a lytic transglycosylase. Importantly, the addition of either DipM or DipM^LytM^ led to a considerable increase in the activity of SdpA against crosslinked PG, while its activity against non-crosslinked PG was only moderately stimulated under these conditions (**Figure 4B,C**). SdpB activity was below the detection limit in our experimental setting, suggesting that this enzyme requires as-yet-unknown additional factors or conditions in order to act. We also tested the effect of DipM on AmiC, assaying the AmiC-mediated cleavage of muropeptides released from PG sacculi by previous muramidase treatment (**Figure S2A**). We found that AmiC had weak basal activity, which was strongly stimulated by DipM or DipM^LytM^ (**Figure S2B**), supporting and extending previous findings based on dye-release assays (Meier et al., 2017). Collectively, our findings demonstrate that DipM not only recruits several autolytic enzymes but also stimulates the activities of SdpA and AmiC, making it the first reported multi-class autolysin activator.

**Figure 4.**
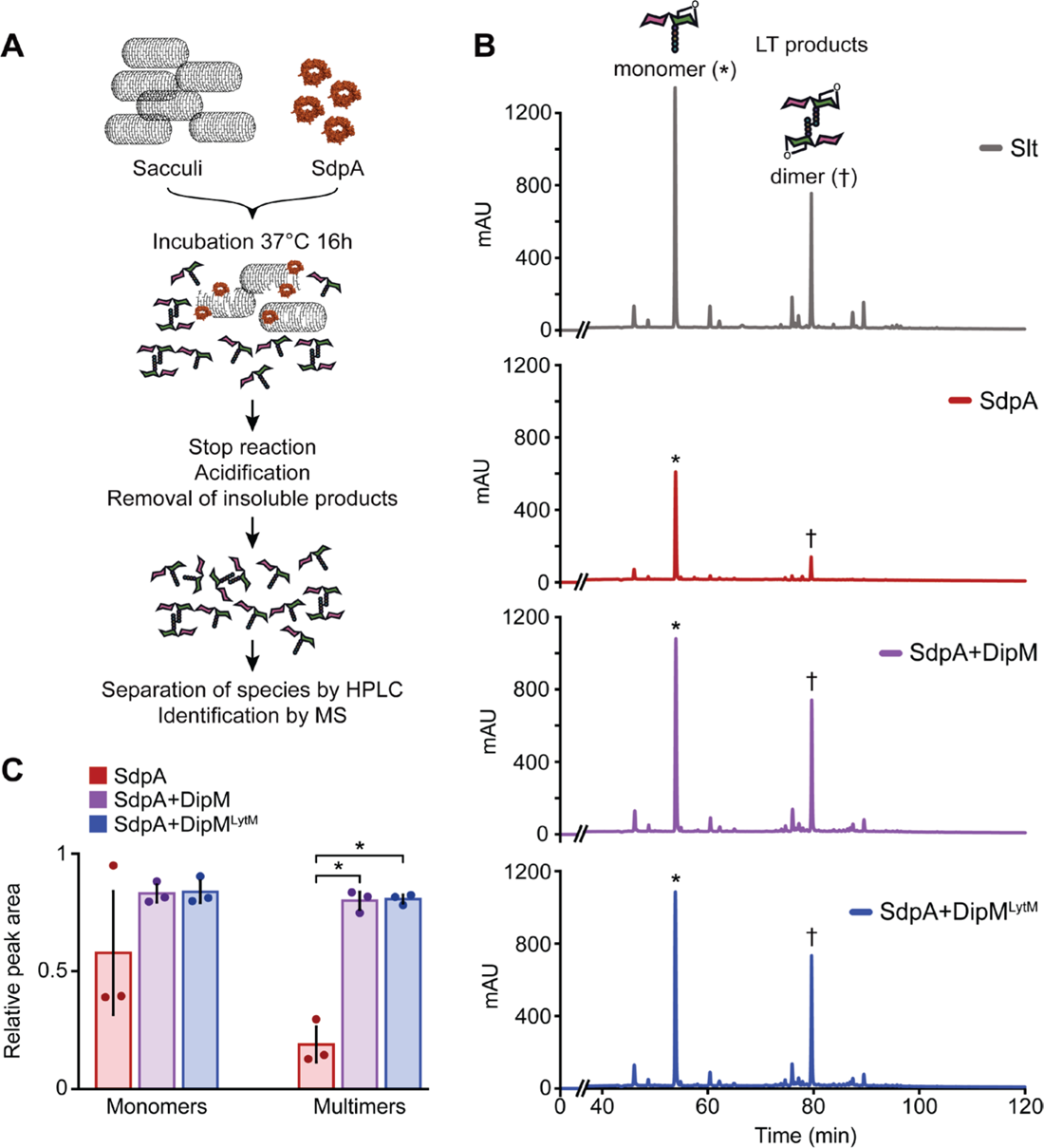
DipM stimulates the lytic transglycosylase activity of SdpA *in vitro*. **(A)** Overview of the procedure used to assess the lytic activity of SdpA. **(B)** HPLC chromatograms showing the muropeptides generated by incubation of sacculi with the indicated protein(s). SdpA and DipM/DipM^LytM^ were used at equimolar ratios. Asterisks mark the peaks of monomeric anhydromuramyl-containing products, daggers those of dimeric anhydromuramyl-containing products. **(C)** Bar chart representing the mean (n=3, individual values are plotted as dots) of the relative area under the peaks of the indicated species under the indicated conditions in panel B. The values have been normalized to those obtained for Slt. Error bars represent the standard deviation and asterisks indicate a significant (*p*<0.001) difference between the means of the conditions.

### The LytM domain of DipM is essential but not sufficient for proper DipM function

Previous work has suggested that the production of LytM domain of DipM was sufficient to complement the Δ*dipM* phenotype (Goley et al., 2010; Möll et al., 2010; Poggio et al., 2010). However, these studies did not provide robust quantifications and employed strains that were constructed by transformation of a Δ*dipM* mutant, which in our hands is phenotypically unstable. To obtain more reliable information about the contributions of the different domains to DipM function, we carefully analyzed the functionality of DipM derivatives that lacked different parts of the N-terminal region including the LysM PG-binding domains (**Figure 5A**). The mutant proteins were tagged with the cyan fluorescent protein sfmTurquoise2^ox^ (Meiresonne et al., 2019) and produced in a conditional *dipM* mutant, whose native DipM protein was depleted right before analysis, avoiding the accumulation of suppressor mutations. Using this system, we found that none of the LysM-less variants was able to restore normal cell division in DipM-depleted cells, whereas a wild-type DipM-sfmTurquoise2^ox^ fusion was fully functional at similar accumulation levels (**Figure B-D**). Notably, the LytM domain alone (DipMΔ35-459) was less active than a variant that additionally contained part of the non-structured region immediately adjacent to the LytM domain (Δ35-390). By contrast, it was more active than a variant comprising the N-terminal non-structured region fused to the LytM domain (DipMΔ123-459), even though the latter unexpectedly showed some accumulation in the mid-cell region (**Figure 5B**). As observed previously (Goley et al., 2010; Möll et al., 2010; Poggio et al., 2010), and in line with the central role of the LytM domain in the activation of DipM target proteins, a variant lacking the C-terminal LytM domain (DipMΔ390-609) was completely non-functional (**Figure S3**). We noticed that the phenotypic differences between the strains were more pronounced in double-concentrated culture medium (data not shown), suggesting that the N-terminal part of DipM may be particularly relevant under conditions of fast growth. Thus, although the function of DipM is largely mediated by the C-terminal LytM domain, the long non-structured regions and the LysM domains contribute to making cell division more robust.

**Figure 5.**
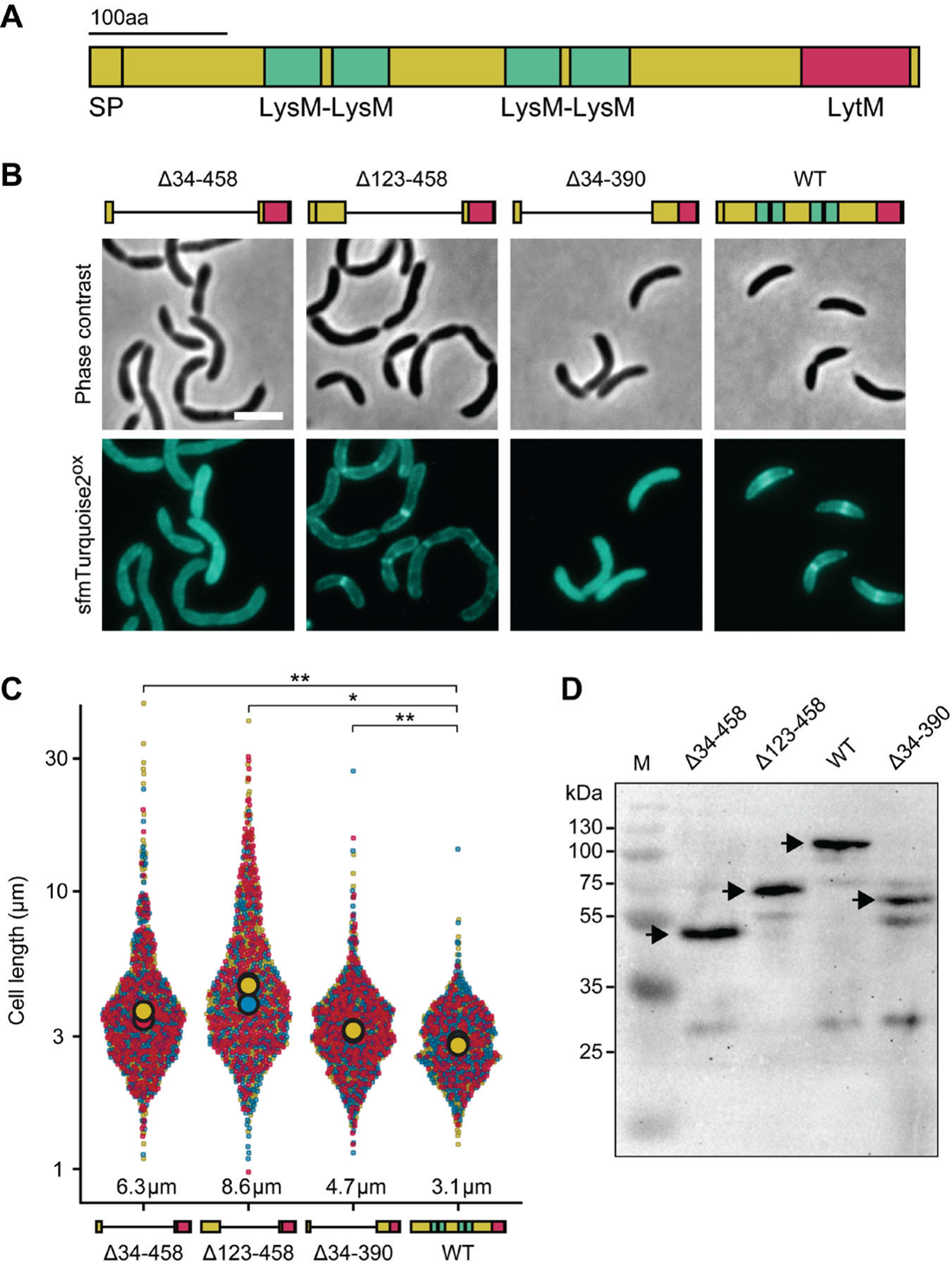
The LytM domain of DipM alone cannot fully complement the loss of the native DipM protein. **(A)** Schematic showing the domain architecture of DipM. The positions of the signal peptide (SP), the four LysM domains and the LytM domain are indicated. Non-structured regions are shown in yellow. **(B)** Localization patterns and functionality of full-length DipM (AI093) or variants lacking parts of the N-terminal region adjacent to the LytM domain (AI091, AI092, AI101). Shown are phase contrast and fluorescence images of cells producing the indicated DipM-sfmTurquoise2^ox^ fusions in place of the wild-type protein in 2xPYE medium. Schematics depicting the domain organization of the different variants are shown on top. **(C)** Superplots showing the distribution of cell lengths in the cultures described in panel B. The data represent the results of three replicates, which are displayed in different colors (red, blue and yellow). The big filled circles indicate the mean cell length obtained for each replicate. Asterisks indicate the statistical significance of differences between the different strains (* p<0.05, ** p<0.005). **(D)** Western blot analysis of the strains analyzed above with an anti-GFP antibody (which also recognizes sfmTurquoise2^ox^). Arrows indicate the expected size of the indicated DipM-sfmTurquoise2^ox^ variants.

### The crystal structure of DipM^LytM^ reveals unique elements compared to EnvC^LytM^

Since the LytM domain is essential for DipM function and sufficient to stimulate SdpA and AmiC activity, we aimed to solve its crystal structure to obtain more insight into its mode of action. For this purpose, we purified a DipM fragment comprising the LytM domain (residues 503-609) and part of the adjacent N-terminal non-structured region (residues 459-502) and solved its molecular structure by X-ray diffraction to 2.25 Å resolution (**Figure 6A** and **Table S1**). The crystals contained four independent chains per asymmetric unit, with clear electron densities for the LytM domain and part of the N-terminal extension (**Figure S4**).

**Figure 6.**
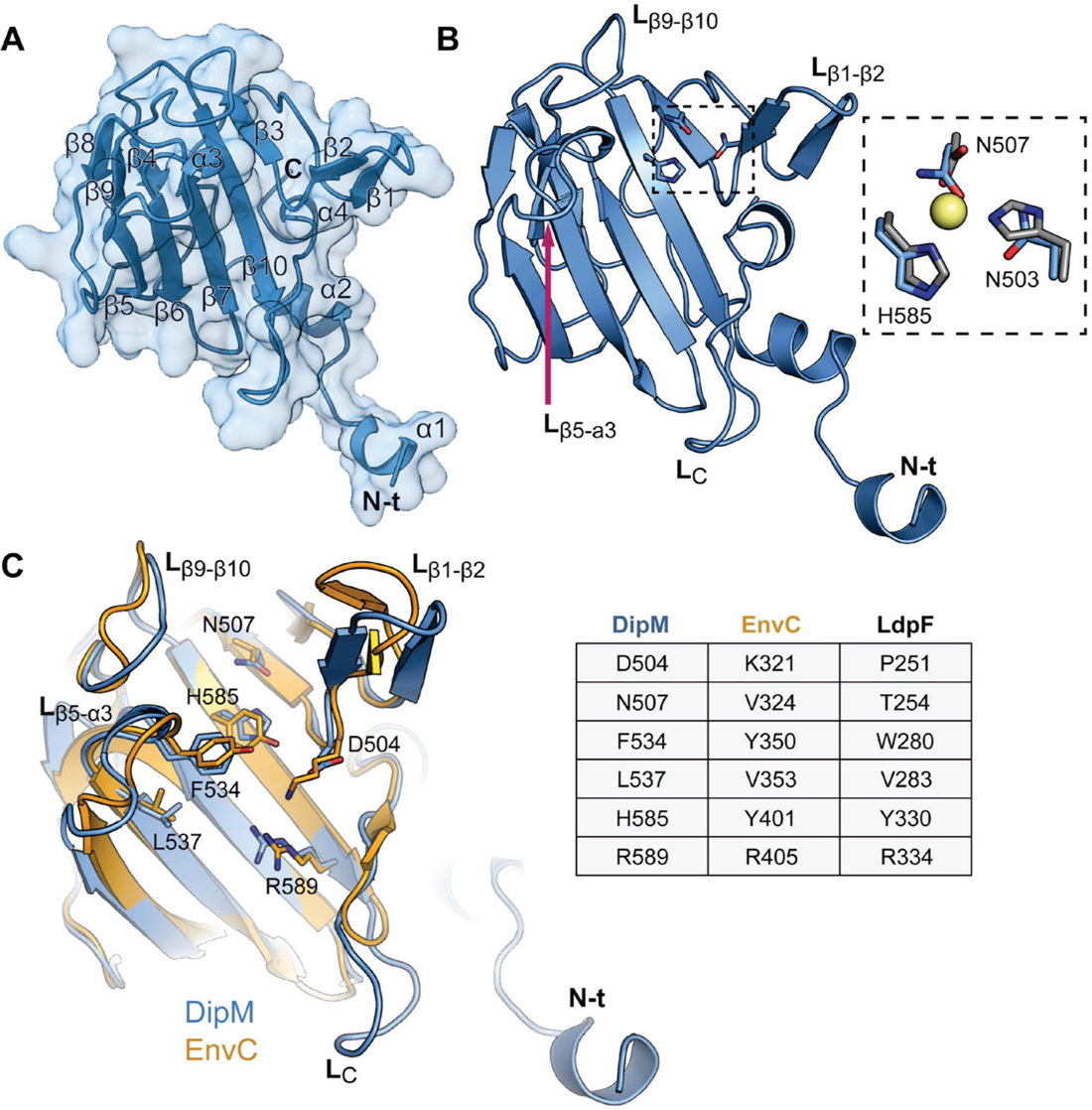
Three-dimensional structure of the LytM domain of DipM. **(A)** Cartoon and surface view of the LytM domain of DipM (DipM^LytM^). Secondary structures are numbered according to their position in the polypeptide chain. **(B)** Cartoon view of DipM^LytM^ in the same orientation as in panel A, with some of the loops labeled. The degenerated active site is highlighted by a frame. A magnification of this region is shown on the right, with the residues of DipM (blue sticks and labeled) superimposed to the metal-binding residues of LytM from *S. aureus* (gray sticks and the catalytic zinc ion shaped as a yellow sphere; PDB: 1QWY). **(C)** Structural alignment of DipM^LytM^ (teal) and *E. coli* EnvC^LytM^ (orange, PDB: 4BH5) (Peters et al., 2013) in cartoon view (rmsd 0.457 Å for 80 C_α_ atoms). Regions showing significant differences are labeled as in panel B. Key residues of EnvC involved in amidase activation as well as the corresponding residues of DipM are depicted as orange and blue sticks, respectively. A comparison of these residues in the LytM factors DipM, EnvC and LdpF is given in the table on the right.

The structure of the LytM domain (DipM^LytM^) is characterized by a core folding motif that comprises a two-layered β-sandwich composed of a central large β-sheet (β3, β5, β6, β7 and β10) and an adjacent smaller β-sheet (β4, β8 and β9) (**Figure 6A**). Beyond this, DipM^LytM^ contains two additional β-strands, forming a β-hairpin (β1-β2) that emanates from the core, a feature also observed for LytM of *S. aureus* and Lysostaphin-like proteins (Bochtler et al., 2004) (**Figure 6A**). As expected for a non-catalytic LytM factor, DipM^LytM^ shows a degenerate catalytic site containing only one of the three metal-coordinating conserved in LytM domains with lytic activity (**Figure 6B**).

The four independent chains in the crystal exhibit a very similar structure, with root-mean-square deviation (rmsd) values for the C_α_ atoms ranging from 0.061 to 0.235 Å. The main differences observed are located in four loops that display conformational plasticity, including L_β1-β2_, L_β5-α3_, L_β9-β10_ and L_C_ (residues 589-597 at the C-terminal end). Differences are also observed in the flexible N-terminal region immediately adjacent to the LytM domain (**Figure S4A,B**). Unless indicated otherwise, chain C will serve as a reference in this study. Overall, the structure of DipM^LytM^ is similar to that of EnvC^LytM^ from *E. coli* (PDB 4BH5,), with an rmsd of 0.457 Å for the 80 superimposed C_α_ atoms (**Figure 6C** and **Figure S4**). However, relevant differences are observed in the before mentioned loops L_β1-β2_, L_β5-α3_, L_β9-β10_ and especially in loop L_C_, which is much longer in DipM than in EnvC and other characterized LytM domain-containing proteins (**Figure 6C** and **Figure S5A**). This extension of L_C_ appears to be conserved only in DipM proteins of the genera *Caulobacter* and *Phenylobacterium* and contains a characteristic KDK motif at its center (**Figure S6**). However, complementation experiments show that this longer loop is not essential for DipM function (**Figure S5B**).

Previous work has shown that the residues of EnvC required for amidase activation are all clustered in and around the larger β-sheet of the LytM domain core fold (Peters et al., 2013). Some of these residues (V353, R405, Y350) are conserved in DipM^LytM^ (L537, R589, F534) and also in LdpF^LytM^ (**Figure 6C**). Others, by contrast, have changed in DipM and adopted different physico-chemical properties, including D504 (K321 in EnvC), N507 (V324 in EnvC) and H585 (Y401 in EnvC) (**Figure 6C**), which may have implications for AmiC recognition (as described below). The positive electrostatic surface potential in this region, which was shown to be essential for amidase activation by EnvC (Peters et al., 2013), is also conserved in DipM, with loop L_c_ providing even more positive charges to the binding pocket (**Figure S7**).

The N-terminal part of the DipM fragment crystallized in this study, ranging from its start to loop β1-β2, is not part of the LytM domain as annotated by the Pfam database (Mistry et al., 2021) (**Figure S8A**), because the underlying hidden Markov model does not include this region and thus predicts the start of the LytM domain at residue Q501 of DipM. However, helix α2 interacts with the C-terminal part of the LytM domain, and the adjacent non-structured region makes close contacts with the LytM core domain. Derivatives of DipM^LytM^ (DipMΔ35-458) lacking parts of this N-terminal region showed significantly reduced protein levels (**Figure S8B**). The strongest effect on DipM accumulation was observed upon the removal of residues 500-479, indicating that this part of the protein may be required for efficient folding or stabilization of the LytM domain. Notably, crystal structures of other LytM domain-containing proteins display comparable non-structured regions with strikingly similar conformations adjacent to the predicted LytM domain (**Figure S8C**). This structural element thus appears to be an integral part of the LytM domain, although its high sequence variability prevents its recognition by a hidden Markov model, based on protein sequence alignments.

### Model for the DipM^LytM^-AmiC interaction in *C. crescentus*

The high structural similarity of the LytM domains of DipM and EnvC prompted us to explore whether partner recognition by DipM could follow the same principles as that by EnvC. Recent work has solved the crystal structure of full-length EnvC in complex with the extra-cellular domain of FtsX (FtsX^ED^) (Cook et al., 2020). In this structure, residues involved in amidase activation by EnvC^LytM^ were found to be occluded by a “restraining arm” formed by an α-helical region adjacent to the LytM domain, suggesting a self-inhibition mechanism (Cook et al., 2020). The interaction between the LytM domain and the restraining arm of EnvC is mediated by hydrophobic, polar and charged residues from both of these protein regions. The determinants in the binding groove that accommodates the restraining arm include residues E317, K321 and Q315 in the β-hairpin motif as well as R340 and E357 in the large β-sheet (**Figure 7B**), matching the residues shown to be involved in amidase activation (Peters et al., 2013). However, the molecular mechanism whereby EnvC stimulates its cognate amidases remains unclear.

**Figure 7.**
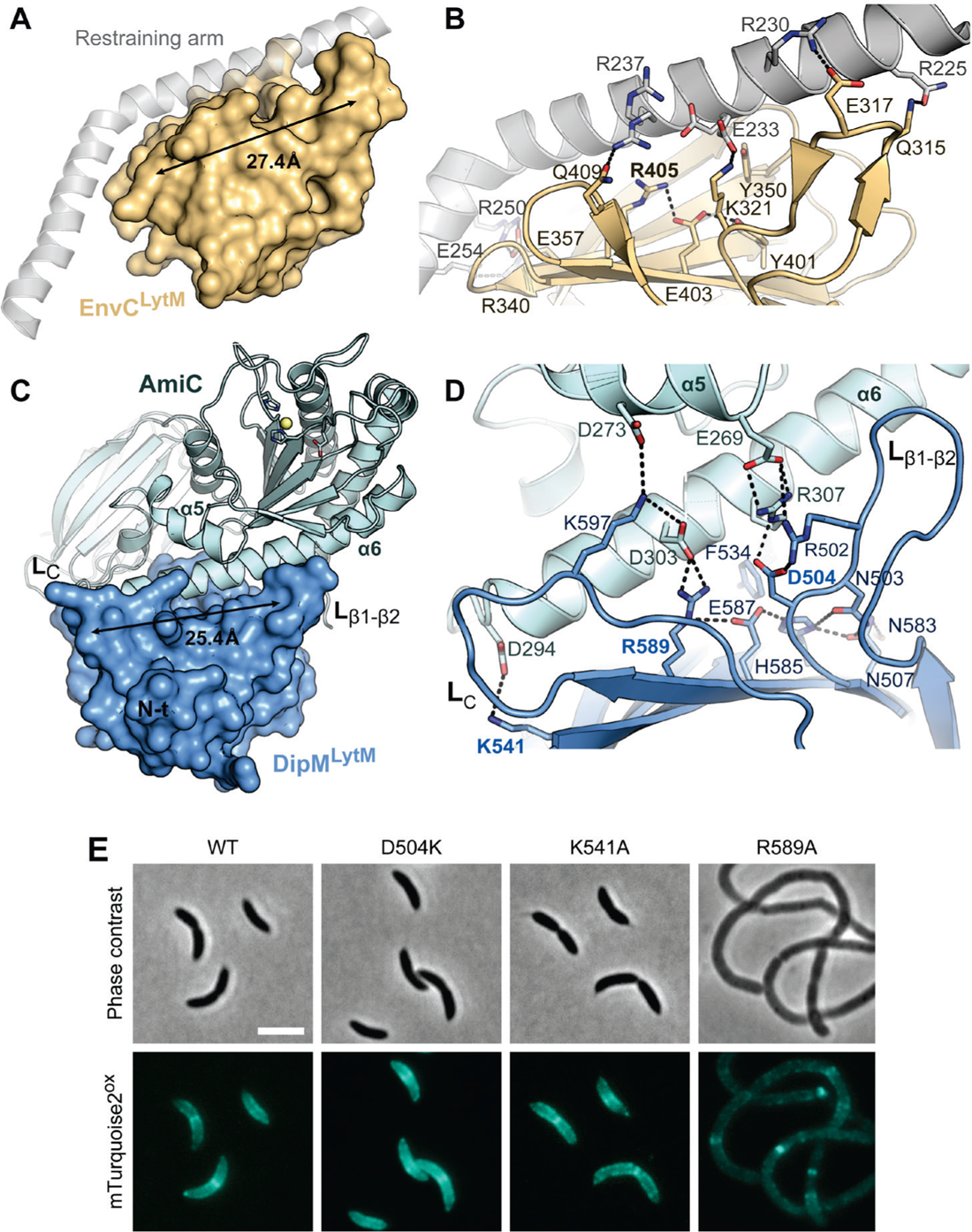
Structural model of the DipM^LytM^-AmiC complex. **(A)** LytM domain of EnvC (EnvC^LytM^) associated with the self-inhibitory restraining arm (gray transparent cartoon; PDB: 6TPI) (Cook et al., 2020). The length of the binding groove is indicated. **(B)** Detailed view of the interface between EnvC^LytM^ and the restraining arm. Hydrogen bonds are displayed as dashed lines. **(C)** Model of the DipM^LytM^-AmiC complex, generated by AlphaFold-Multimer (Evans et al., 2022). The crystal structure of DipM^LytM^ is shown in surface representation (dark blue), the modeled structure of AmiC from *C. crescentus* in cartoon view (light blue). The length of the predicted binding groove is indicated. **(D)** Detailed view of the predicted interface between DipM^LytM^ and helices α5/α6 of AmiC. Residues involved in the interaction are shown in stick representation. Relevant hydrogen bonds are shown as dotted lines. **(I)** Functionality of DipM variants with exchanges in the predicted AmiC binding pocket. Shown are phase contrast and fluorescence micrographs of cells producing wild-type (WT) DipM-sfmTurquoise2^ox^ or the indicated mutant derivatives (AI093, AI129, AI132, AI133). Scale bar: 3 µm.

Interestingly, DipM^LytM^ shows a conspicuous groove (9.5 Å x 25.4 Å) whose dimensions are similar to that of the EnvC binding groove (10 Å x 27.4 Å) (**Figure 7A** and **Figure S9A,B**). To obtain more insight into the potential mode of action of DipM, we used AlphaFold-Multimer (Evans et al., 2022) to generate a structural model of DipM^LytM^, which agreed very well with the crystal structure solved in this study (**Figure S10A**). We then went on to generate a model for the DipM^LytM^-AmiC complex in *C. crescentus*. The five top-ranking models generated by AlphaFold-Multimer were highly similar to each other (**Figure S10B,C**). Interestingly, they predicted that the interaction of AmiC with DipM^LytM^ was mediated by a long α-helix (α6) from AmiC that was inserted into the groove of DipM^LytM^ (**Figure 7C**), recapitulating the arrangement of the restraining arm in the binding groove of EnvC^LytM^ (**Figure 7A**).

As for EnvC^LytM^, the groove of DipM^LytM^ is limited by loops L_β1-β2_, L_β5-α3_, L_β9-β10_ and L_C_ (**Figure S10C,D**) and covered with hydrophobic residues. In the predicted complex, residues in loops L_β1-β2_ and L_C_ closely interact with helix α6 of AmiC (**Figure 7C,D**). Additional interactions are made with helix α5 of AmiC (**Figure 7C**). It is conceivable that complex formation induces conformational changes in AmiC that activate its amidase activity. A detailed analysis of the predicted DipM^LytM^-AmiC complex identified residues that could potentially be critical for amidase recognition and/or activation. Conserved residue R589 may promote these processes by forming a salt bridge with D303 in helix α6 of AmiC (**Figure 7D**). Its proper orientation is likely ensured by an extensive hydrogen bonding network involving residues E587, H585, N503, N507 and E587. Loop L_C_ may contribute to AmiC binding through residue K597, which is predicted to interact with both residue D273 in helix α5 and residue D303 in helix α6 of AmiC (**Figure 7D**). Similarly, residues R502 and D504 in loop L_β1-β2_ form salt bridges with residues E269 and R307, respectively, in these to helices (**Figure 7D**).

To test the structural model, we aimed to determine whether modifications in the putative AmiC-binding interface affected the function of DipM. To this end, we exchanged residues D504, K541 or R589, which are located in different regions of the groove, and analyzed the functionality of the mutant proteins in a complementation assay (**Figure 7E**). Two of the variants (D504K and K541A) did not show any obvious defects, potentially suggesting redundancy in the interaction determinants. The R589A exchange, by contrast, led to a complete loss of DipM activity, producing a phenotype similar to that observed for the Δ*dipM* mutant. This finding supports the notion that the groove in the LytM domain of DipM serves as a docking site for AmiC and potentially other DipM targets. Moreover, it suggests that EnvC could use a similar mechanism to achieve amidase activation once the FtsEX complex triggers the dissociation of the auto-inhibitory restraining arm from its LytM domain.

### Independence of protein localization and activation in the FtsN-DipM-SdpA/B pathway

Previous work has shown that DipM is required to localize the lytic transglycosylases SdpA and SdpB to the mid-cell region (Zielińska et al., 2017). Since DipM interacts with SdpA and stimulates its activity, we tested whether the recruitment of SdpA and SdpB depended on direct protein-protein interactions with DipM. To this end, we determined the localization patterns of the two proteins in a strain that produced a truncated variant of DipM lacking the LysM domains (DipMΔ36-458), which supported cell division even though it was evenly distributed within the cell (compare **Figure 5**). In this background, both SdpA and SdpB were still recruited to the cell center (**Figure 8**), whereas they failed to do so in a Δ*dipM* mutant (Zielińska et al., 2017). Thus, DipM does not recruit its two targets through direct physical interaction but rather indirectly through the regulatory activity of its LytM domain during cell division.

**Figure 8.**
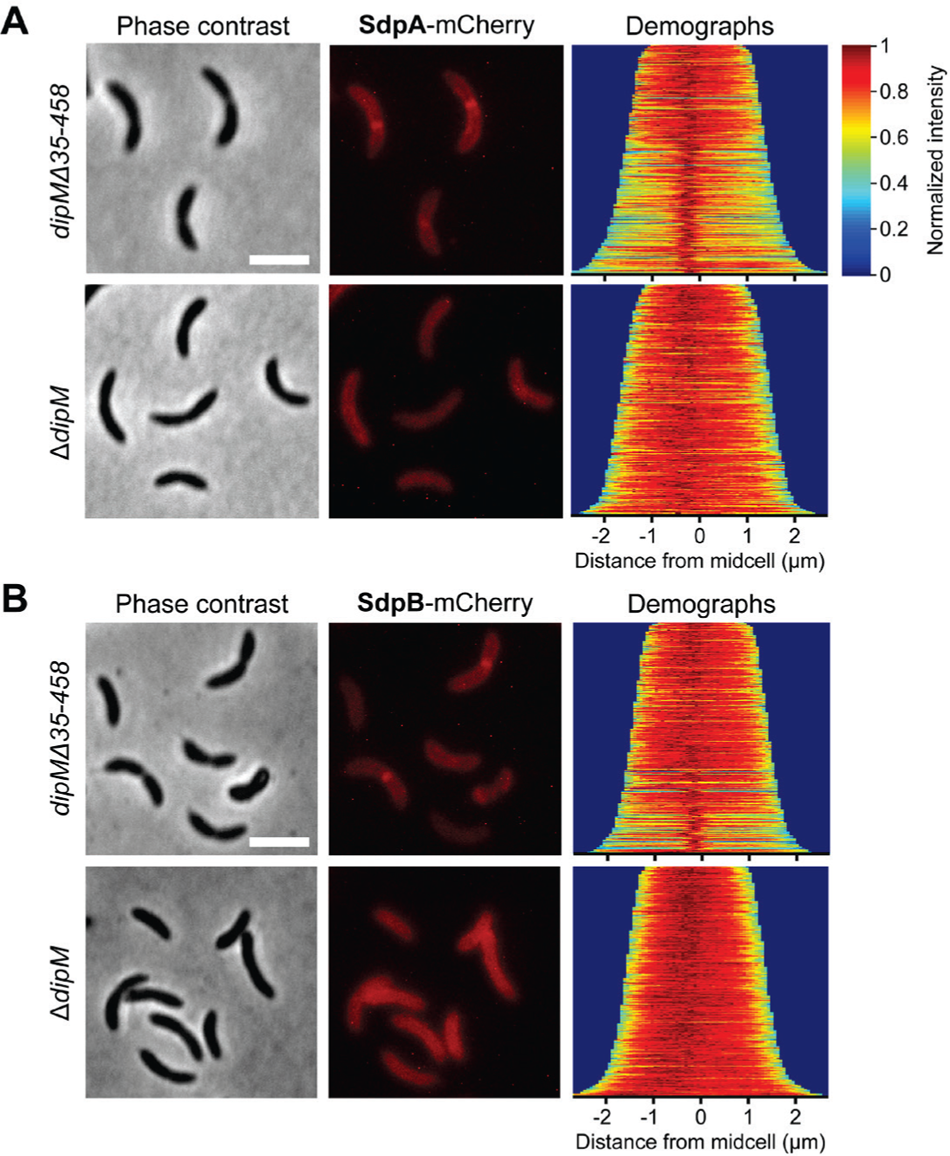
Localization dependencies in the FtsN-DipM-SdpA/B regulatory pathway. **(A)** Localization of SdpA-mCherry in the Δ*dipM* (MAB203) and *dipMΔ35-458* (AI113) backgrounds. Cells were grown in PYE medium and induced to produce the fluorescent protein fusion by addition of 0.3% xylose. After two more hours of incubation, the population was synchronized, transferred to PYE medium containing 0.3% xylose and allowed to grow for 75 min prior to imaging by phase contrast and fluorescence microscopy. The demographs show the fluorescence profiles of a representative subpopulation of cells ordered according to cell length and stacked on top of each other (n>300 each). **(B)** Localization of SdpB-mCherry in the Δ*dipM* (MAB308) and *dipMΔ35-458* (AI114) backgrounds. Cells were analyzed as described in panel A.

Prompted by these results, we also investigated the requirements for the recruitment of DipM. DipM directly associated with the SPOR domain of FtsN *in vitro* (**Figure 3E,F**) and no longer localized to the cell center upon FtsN depletion (Goley et al., 2010), suggesting a role of direct protein-protein interactions in the recruitment process. However, DipM was still able to accumulate at the division site in cells whose native FtsN protein was replaced by a truncated variant lacking the SPOR domain, which fails to condense in the mid-cell region (Möll and Thanbichler, 2009) (**Figure S11**). This finding demonstrates that the decisive role of FtsN in DipM localization is also indirect and likely based on FtsN-dependent PG remodeling at the division site (Du and Lutkenhaus, 2017; Müller et al., 2007; Weiss, 2015), which may induce specific modifications in the constricting PG layer that are then recognized by the LysM domains of DipM. Thus, physical interactions are not the main drivers of protein localization in the FtsN-DipM-SdpA/B regulatory pathway, even though its components do interact with each other.

### SdpA and SdpB affect the single-molecule dynamics of DipM

The LytM domain of DipM alone can mediate cell division, although it is evenly distributed within the cell (**Figure 5**), suggesting that transient interactions with its regulatory targets at the division site are sufficient for close-to-normal activity. To investigate the regulatory interplay between DipM and its target proteins in more detail, we set out to perform single-molecule tracking studies of DipM. To this end, the full-length protein, a C-terminally truncated variant lacking the LytM domain (DipM^ΔLytM^) and the LytM alone (DipM^LytM^) were fused to sfmTurquoise2^ox^ and produced in both a wild-type or Δ*sdpAB* background (**Figure S12**). Subsequently, we followed the motion of single molecules in synchronized live cells harvested shortly after the onset of cell constriction, a cell cycle stage when DipM is usually localized at the cell division site (Goley et al., 2010; Möll et al., 2010; Poggio et al., 2010). An analysis of the single-molecule tracks obtained revealed that overall mobility of DipM was very low, with most molecules showing slow, confined motion in the central region of the cell (**Figure 9A**). In each of the cells imaged, a smaller fraction of slow-moving molecules was also observed at one of the cell poles. Our experimental setting did not allow us to discriminate between the old and new cell pole. However, DipM is known to localize to the old pole early in the cell cycle to support stalk formation (Billini et al., 2019), suggesting that these signals reflect the old-pole-associated DipM population. The distribution of step sizes in the single-particle tracks indicates the existence of two distinct diffusion regimes, with a slow-moving (static) and a fast-moving (mobile) population (**Figure S13**). For full-length DipM, ∼75% of the molecules were static, consistent with previous fluorescence-recovery-after-photobleaching data (Poggio et al., 2010), whereas the remaining 25% were mobile, with an ∼10-fold higher diffusion rate (**Figure 9B,C**).

**Figure 9.**
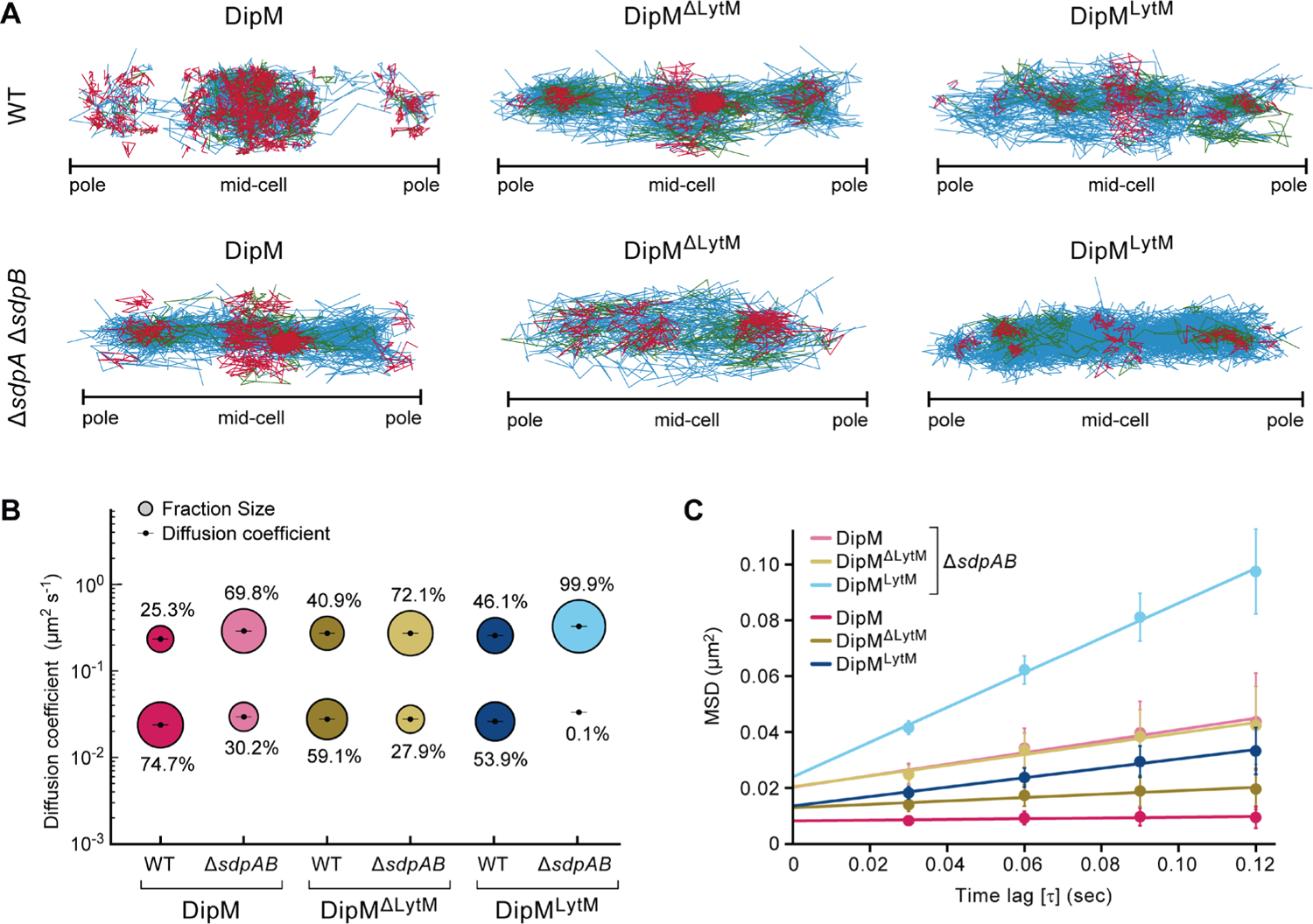
SdpA and SdpB affect the single-molecule behavior of DipM. **(A)** Confinement maps showing single-molecule tracks of DipM, DipM_1-390_ (DipM^ΔLytM^) or DipM_Δ35-458_ (DipM^LytM^) fused to sfmTurquoise2^ox^ in the wild-type (AI063, AI098, AI112) and Δ*sdpAB* (AI121, AI122, AI126) backgrounds. Cells producing the indicated fluorescent protein fusions were synchronized and allowed to grow for another 75 min prior to imaging. Confined tracks are shown in red, non-confined tracks in blue, and tracks showing both behaviors in green. Note that, due to technical limitations, it was not possible to determine the orientation of the cells and, thus, distinguish between the stalked (old) and non-stalked (new) pole during imaging. Data represent the results of two biological replicates. **(B)** Bubble plots showing the proportions of static and mobile molecules for the indicated DipM-sfmTurquoise2^ox^ fusions in the wild-type and Δ*sdpAB* backgrounds. **(B)** Mean-squared-displacement (MSD) analysis of the indicated DipM-sfmTurquoise2^ox^ fusions in the wild-type and Δ*sdpAB* backgrounds. The chart shows the MSD (± SD) values for the first four time lags (τ) from all tracks containing more than four points and linear fits of the data for each of the indicated conditions.

The two truncated DipM variants were more mobile than the full-length protein, even though both DipM^ΔLytM^ and, to a lesser degree, DipM^LytM^ still showed an appreciable number of confined tracks at the division site (**Figure 9A**). For DipM^ΔLytM^, the proportion of static molecules was only slightly lower (∼60%) (**Figure 9B,C**), in line with the notion that the LysM domains in the N-terminal region of DipM tether the protein to nascent PG at the cell division site and the stalk base. Surprisingly, however, DipM^LytM^ also showed a large fraction (∼54%) of static molecules. This finding indicates that the LytM domain closely interacts with components of the cell division apparatus, explaining why it largely retained its functionality, even though the population as a whole no longer accumulated at the division site (**Figure 5**).

Importantly, the mobility of DipM and its variants increased considerably in cells lacking the two lytic transglycosylases SdpA and SdpB, accompanied by a decrease in the fraction of confined tracks at the cell center or poles (**Figure 9A**). Under these conditions, the proportion of static full-length DipM and DipM^ΔLytM^ molecules decreased to ∼30% (**Figure 9B,C**). By contrast, confined motion became essentially undetectable for DipM^LytM^, indicating that the interaction of the LytM domain with the division apparatus is mediated by either or both of the two DipM target proteins. These results reveal the existence of a self-reinforcing cycle in which the DipM activity stimulates the accumulation of SdpA and SdpB at the cell division site, which then in turn promote the recruitment of DipM to the mid-cell region, either through direct physical interaction with its LytM domain or through modifications to the PG layer that are recognized by its PG-binding LysM domains.

## Discussion

Proteins with catalytically inactive C-terminal LytM domains, termed LytM factors, are found throughout the phylum Proteobacteria and also in some cyanobacteria. Some of their gammaproteobacterial representatives, such as EnvC, contain coiled-coil regions and mediate the interaction between the FtsEX complex and amidases (Möll et al., 2014; Uehara et al., 2009; Uehara et al., 2010; Yakhnina et al., 2015; Yang et al., 2012). The LytM factor LdpF from *C. crescentus* resembles these proteins in its domain architecture, and our Co-IP data indicate that it may indeed serve to physically connect FtsEX with the amidase homolog AmiC (**Figure 1E,F**). However, it has been shown that LdpF does not play a role in AmiC activation (Meier et al., 2017), although it is required for its recruitment to the mid-cell region (Zielińska et al., 2017). Instead, another LytM factor, DipM, has been adopted as an additional regulator to stimulate AmiC activity in the *C. crescentus* system (Meier et al., 2017) (**Figure S2B**). It is possible that LdpF served as an amidase activator in the past but lost this activity during the course of evolution to allow for more complexity in the control of amidase activity. This hypothesis is in agreement with the idea, born from studies in *E. coli*, that amidases and their activators are recruited independently of each other, creating a fail-safe system that minimizes aberrant PG degradation during cell division (Peters et al., 2011). It remains to be clarified whether this concept also applies to other alphaproteobacteria and how tight amidase regulation is achieved in species such as *H. neptunium*, which possess only a single catalytically inactive LytM factor (Cserti et al., 2017).

Other gammaproteobacterial LytM factors, such as NlpD and ActS from *E. coli*, contain PG-binding LysM domains instead of coiled-coil regions and mediate amidase activation in an FtsEX-independent manner (Gurnani Serrano et al., 2021; Tsang et al., 2017; Uehara et al., 2009; Yang et al., 2018). *C. crescentus* DipM bears resemblance to these proteins, although its LysM domains are organized into two tandems, an arrangement not found in gammaproteobacteria and other alphaproteobacteria. Given its role in amidase activation and its apparent lack of interaction with FtsEX (**Figure 1A,F**), DipM could be considered an NlpD homolog. However, while the role of NlpD in *E. coli* is likely limited to connecting AmiC with the Tol/Pal system (Tsang et al., 2017), DipM interacts with FtsN and at least four different autolysins, thus exhibiting an unprecedented regulatory complexity (**Figure 1G**). We show that DipM not only stimulates the lytic activity of AmiC (**Figure S2B**) but also that of SdpA (**Figure 4B,C**). It thus appears that, at least in the Alphaproteobacteria, LytM factors can be multi-class autolysin activators, controlling both amidases and lytic transglycosylases. Importantly, while self-regulation has been observed for lytic transglycosylases of family 1E (Domínguez-Gil et al., 2016), this is, to our knowledge, the first reported case of lytic transglycosylase activation through protein-protein interaction. Notably, a recent study in *E. coli* has also reported a novel lipoprotein, called NlpI, that can interact with several endopeptidases and connect them to other components of the peptidoglycan biosynthetic machinery (Banzhaf et al., 2020). Thus, factors that orchestrate the activities of multiple autolysins might be widespread in bacteria.

The ability of DipM^LytM^ to activate two different classes of enzymes opens the question of whether the binding of its interaction partners is mediated by the same or rather by different interfaces. The results of our BLI-based competition assays support the hypothesis that the binding interfaces could at least overlap, because AmiC and SdpA appear to compete for DipM (**Figure S1**). In line with this notion, the R489A exchange, which we predicted to affect AmiC binding, completely abolished DipM function (**Figure 7E**), suggesting that modifications in this interface affect multiple interactions and not only the one with AmiC. We expect that future structural analysis can clarify the precise mechanisms underlying the interactions and stimulatory effects mediated by DipM.

Based on the results of this and previous studies, we hypothesize that DipM functionally connects two protein-recruitment and enzyme-activation cycles in *C. crescentus*: one involving FtsN, AmiC and other late divisome components and a second one involving the two SLTs SdpA and SdpB (**Figure 10**). Since DipM is the activator of AmiC, it is responsible for the generation of naked PG, which is in turn recognized by SPOR domain-containing proteins such as FtsN (Alcorlo et al., 2019; Yahashiri et al., 2015, 2017). It has been proposed that this synergistic interplay between amidases and the SPOR domain is the main driving force for the recruitment of FtsN to the division site, which in *E. coli* acts as trigger for the activation of late divisome components and, thus, septum formation (Gerding et al., 2009; Lutkenhaus, 2009). We have previously shown that DipM requires FtsN for its recruitment to the mid-cell region (Zielińska et al., 2017). Interestingly, this dependency does not rely on a physical interaction of DipM with the periplasmic region of FtsN (**Figure S11**), indicating that it is rather based on the regulatory activity of FtsN. We hypothesize that FtsN affects the localization of DipM indirectly by stimulating the synthesis of septal PG, thereby generating structural cues in the PG layer that are recognized by the PG-binding LysM domains of DipM. These interactions give rise to a self-reinforcing cycle that promotes the progressive accumulation of DipM and FtsN at the division site, in a process that depends on amidase activity and, in turn, gradually increases AmiC activity in the mid-cell region as cell constriction proceeds.

**Figure 10.**
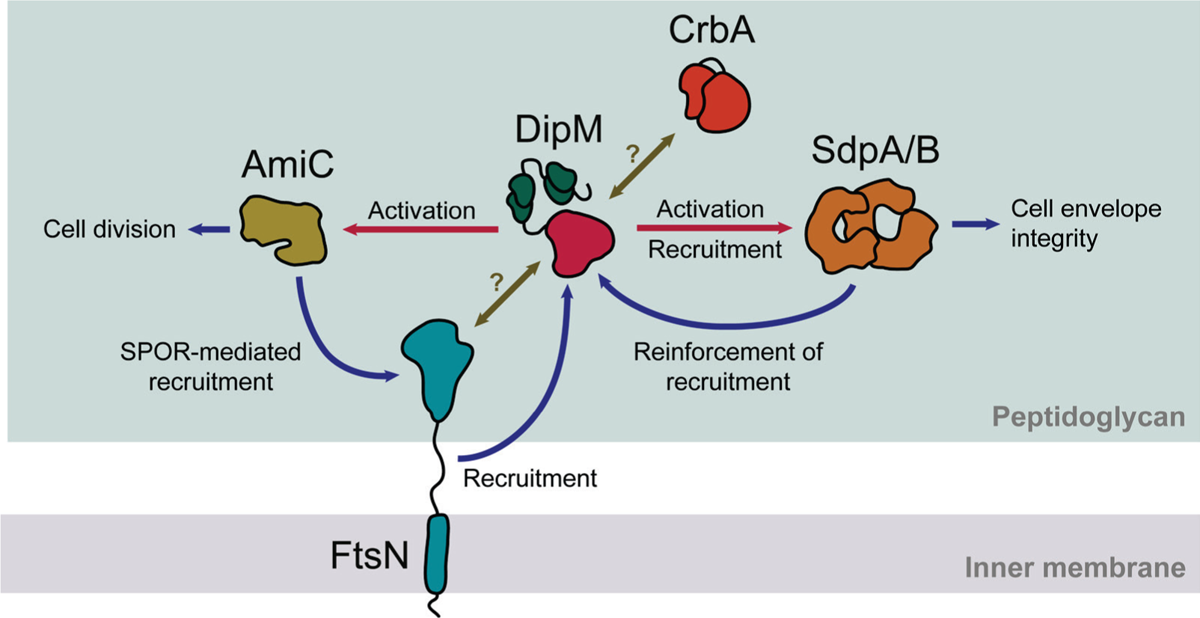
DipM mediates two self-reinforcing cycles that promote peptidoglycan remodeling during cell division. Model for the proposed function of DipM in linking the accumulation and/or activity of the PG biosynthesis regulator FtsN, the amidase AmiC, and the lytic transglycosylases SdpA and SdpB. The effects that the different proteins exert on each other are indicated by arrows (red: enzyme activation, blue: recruitment and other stimulatory effects, olive: unknown).

Our results further show that SdpA and SdpB have a profound effect on the behavior of both the N-terminal LysM domains and the C-terminal LytM domain of DipM. The presence of these two lytic transglycosylases promotes the immobilization of DipM molecules at the division site and at polar locations (most likely the stalked pole) (**Figure 9**). Conversely, the LytM domain of DipM is necessary for the recruitment of SdpA and SdpB to the mid-cell region, possibly because of its stimulatory effect on PG remodeling at this location. Thus, while the regulatory activity of DipM mediates the positioning of SdpA and SdpB, the two enzymes in turn stabilize DipM in the mid-cell region, creating another self-reinforcing cycle that leads to a progressive increase in lytic transglycosylase activity at the division site.

Together, these two self-reinforcing cycles enable DipM to connect the recruitment and/or activation of lytic transglycosylases to late-divisome assembly and cell division. Notably, previous work has implicated SdpA and SdpB in cell envelope integrity (Zielińska et al., 2017), and studies of their *E. coli* homolog Slt70 suggest a role for lytic transglycosylases in the degradation of excess, non-crosslinked PG (Cho et al., 2014). In support of the latter point, we have observed a higher activity of SdpA against non-crosslinked than crosslinked PG *in vitro*. Considering that *C. crescentus* mostly grows in length through divisome-dependent medial PG incorporation (Aaron et al., 2007), DipM may support this mode of growth by promoting “PG quality control” through SdpA and SdpB.

Collectively, the finding that DipM acts as a multi-class autolysin activator furthers our understanding of the functions that catalytically inactive LytM factors can adopt. Although DipM may be a specific invention of the alphaproteobacterial lineage, it is likely that other bacteria have evolved similar systems to coordinate multiple autolysins with a single regulatory protein. The characterization of such multi-enzyme regulators, whose malfunction compromises several cell wall-related processes at once, will not only improve our understanding of PG remodeling but also reveal promising new targets for the development of antibacterial drugs.

## Materials and Methods

### Growth conditions

All *C. crescentus* strains used in this study are derivatives of the synchronizable wild-type strain NA1000 (CB15N) (Evinger and Agabian, 1977). *C. crescentus* cells were grown aerobically at 28 °C in peptone-yeast extract (PYE) medium (Poindexter, 1964), double-concentrated PYE (2xPYE) medium or M2G medium (Ely, 1991), supplemented when necessary with antibiotics at the following concentrations (µg ml^-1^; liquid/solid medium): kanamycin (5/25), gentamicin (0.5/5). To induce the *xylX* (Meisenzahl et al., 1997) or *vanA* (Thanbichler et al., 2007) promoters, media were supplemented with 0.3% D-xylose or 0.5 mM sodium vanillate, respectively. For microscopic analysis, cells were grown to exponential phase and induced for 2-3 h prior to imaging, unless indicated otherwise. *C. crescentus* swarmer cells were isolated by density gradient centrifugation using Percoll (Tsai and Alley, 2001). Plasmids were introduced by electroporation (Ely, 1991) or conjugation, as described previously for *H. neptunium* (Jung et al., 2015). *E. coli* TOP10 (ThermoFisher Scientific) was used for general cloning purposes, *E. coli* WM3064 (W. Metcalf, unpublished) as donor strain for conjugation and *E. coli* Rosetta(DE3)pLysS (Novagen) as host for recombinant protein production. *E. coli* derivatives were grown aerobically at 37 °C in Luria-Bertani medium (LB), supplemented when required with antibiotics at the following concentrations (µg ml^-1^; liquid/solid medium): gentamicin (15/20), kanamycin (30/50), ampicillin (200/200), chloramphenicol (20/30). Cultures of *E. coli* WM3064 were supplemented with 300 µM 2,6-diaminopimelic acid (DAP).

### Plasmid and strain construction

Details on the strains, plasmids and oligonucleotides used in this study are given in **Supplementary Tables 2, 3 and 4**, respectively. Plasmids were designed with the help of SnapGene (GLS Biotech, USA). All constructs were verified by DNA sequencing. Plasmids bearing inducible genes were integrated ectopically at the chromosomal *xylX* (P*_xyl_*) or *vanA* (P*_van_*) loci of *C. crescentus* by single-homologous recombination (Thanbichler et al., 2007). The truncation or in-frame deletion of genes was carried out using a two-step procedure based on the counterselectable *sacB* marker (Thanbichler and Shapiro, 2006). Proper plasmid integration or gene deletion were verified by colony PCR. The correct truncation of endogenous genes was further confirmed by sequencing of the target loci.

### DipM complementation assays

For complementation experiments, strains expressing *dipM-flag* from the P*_xyl_* promoter and mutant versions of *dipM* from the P*_van_* promoter were inoculated from cryo-stocks into liquid PYE medium supplemented with 0.3% xylose and grown overnight. The next day, cells were collected by centrifugation for 1 min at 8000 ×g and washed in 2xPYE medium without any supplements. They were then used to inoculate two new cultures per strain, one in 2xPYE medium containing 0.5 mM sodium vanillate and one in medium without inducer, to a starting OD_600_ of 1.5 x 10^-4^. After 18-20 h of growth, the cells were imaged, and samples were taken for Western blot analysis when required. The non-induced cells served as controls to verify the depletion of DipM, as reflected by filamentation and the formation of membrane blebs (Goley et al., 2010; Möll et al., 2010; Poggio et al., 2010).

### Immunoblot analysis

Immunoblot analysis was conducted as described previously (Thanbichler and Shapiro, 2006), using anti-GFP polyclonal antibodies (Sigma, Germany). Goat anti-rabit immunoglobulin G conjugated with horse-radish peroxidase (Perkin Elmer, USA) was used as secondary antibody. Immunocomplexes were detected with the Western Lightning Plus-ECL chemiluminescence reagent (Perkin Elmer, USA). The signals were recorded with a ChemiDoc MP imaging system (BioRad, Germany) and analyzed using the Image Lab software (BioRad, Germany). For statistical tests, Microsoft Excel (Microsoft) was used.

### Widefield fluorescence imaging

Cells were grown to exponential phase (unless indicated otherwise) and spotted on 1% agarose pads. Images were taken with an Axio Observer.Z1 microscope (Zeiss, Germany) equipped with a Plan Apochromat 100x/1.45 Oil DIC and a Plan Apochromat 100x/1.4 Oil Ph3 phase contrast objective, an ET-mCherry and ET-CFP filter set (Chroma, USA) and a pco.edge 4.2 sCMOS camera (PCO, Germany). Images were recorded with VisiView 3.3.0.6 (Visitron Systems, Germany) and processed with Metamorph 7.7.5 (Universal Imaging Group, USA), Fiji (Schindelin et al., 2012), Adobe Photoshop CS6 and Adobe Illustrator CS6 (Adobe Systems, USA). Cell length measurements were performed with the ObjectJ plugin (Norbert Vischer; available at https://sils.fnwi.uva.nl/bcb/objectj) of Fiji. Demographs were generated using BacStalk (Hartmann et al., 2018). SuperPlots (Lord et al., 2020) were used to visualize cell length distributions and to evaluate the statistical significance of differences between multiple distributions, employing the PlotsOfData web app (Postma and Goedhart, 2019).

### Single-particle tracking and diffusion analysis

Cells of the indicated strains were cultivated overnight in PYE medium, transferred into fresh medium and grown to exponential phase prior to synchronization. The isolated swarmer cells were transferred into pre-warmed M2G liquid medium supplemented with 0.3% D-xylose and grown for 90 min before imaging by slimfield microscopy. In this approach, the back aperture of the objective is underfilled by reduction of the width of the laser beam, generating an area of about 10 μm in diameter with high light intensity that allows the visualization of single fluorescent protein molecules at very high acquisition rates. The single-molecule level was reached by bleaching of most molecules in the cell for 100 to 1,000 frames, followed by tracking of the remaining and newly synthesized molecules for ∼3,000 frames. Images were taken at 30 ms intervals using an Olympus IX-71 microscope equipped with a UAPON 100x/ NA 1.49 TIRF objective, a back-illuminated electron-multiplying charge-coupled device (EMCCD) iXon Ultra camera (Andor Solis, USA) in stream acquisition mode, and a LuxX 457-100 (457 nm, 100 mW) light-emitting diode laser (Omicron-Laserage Laserprodukte GmbH, Germany) as an excitation light source. The laser beam was focused onto the back focal plane and operated during image acquisition with up to 2 mW (60 W/cm^2^ at the image plane). Andor Solis 4.21 software was used for camera control and stream acquisition. Prior to analysis, frames recorded before reaching the single-molecule level were removed from the streams, using photobleaching curves as a reference. Subsequently, the streams were cropped to an equal length of 2,000 frames and the proper pixel size (100 nm) and time increment were set in the imaging metadata using Fiji (Schindelin et al., 2012). Single particles were tracked with u-track 2.2 (Jaqaman et al., 2008). Trajectories were only considered for further statistical analysis if they had a length of at least five steps. Data analysis was performed using SMTracker 2.0 (Oviedo-Bocanegra et al., 2021). The diffusive behavior of the proteins investigated was analyzed in two ways. Mean-squared-displacement (MSD)-versus-time-lag curves were calculated to provide an estimate of the diffusion coefficient and clarify the kind of motion exhibited (e.g. diffusive, subdiffusive or directed). In addition, we determined the frame-to-frame displacement of all molecules in x and the y direction and fitted the resulting distributions to a two-population Gaussian mixture model to determine the proportions of mobile and static molecules in each condition (Oviedo-Bocanegra et al., 2021).

### Protein purification

#### Full-length DipM

To purify MAS-DipM(26-609)-His_6_, cells of *E. coli* AM201 (Möll et al., 2010) were grown in LB medium at 37°C. At an OD_600_ of 1, protein overproduction was induced by the addition of 1 mM isopropyl-β-D-thiogalactopyranoside (IPTG) and cultivation was continued for 3 h. The cells were harvested by centrifugation, washed with buffer BZ3 (50 mM Tris, 300 mM NaCl, 10% v/v glycerol, 20 mM imidazole, adjusted to pH 8 with HCl) and stored at −80°C until further use. After thawing, they were resuspended in buffer BZ3 (2 ml per 1 g of pellet) supplemented with DNase I (10 µg/ml) and PMSF (100 µg/ml). After three passages through a French press at 16,000 psi, cell debris was removed by centrifugation at 30,000 ×g for 30 min.

The supernatant was then applied onto a 5 ml HisTrap HP column (GE Healthcare, USA) equilibrated with buffer BZ3. The column was washed with 10 column volumes (CV) of buffer BZ3, and the protein was eluted with a linear imidazole gradient (20-250 mM in buffer BZ3) at a flow rate of 1 ml/min. Fractions containing the protein at high purity were pooled and dialyzed against buffer B6 (50 mM HEPES, 50 mM NaCl, 5 mM MgCl_2_, 0.5 mM EDTA, 10% v/v glycerol, adjusted to pH 7.2 with NaOH). Afterwards, the protein was concentrated in a centrifugal filter device (Amicon, USA), aliquoted, snap-frozen in liquid nitrogen and stored at −80°C.

For use in lytic transglycosylase activity assays, MAS-DipM(26-609)-His_6_ was subjected to an additional chromatographic step added after Ni-NTA affinity purification. To this end, the protein was dialyzed against buffer LS (50 mM Tris, 50 mM NaCl, adjusted to pH 8 with HCl) and applied to an HiTrap SP cation exchange column (GE Healthcare, USA) equilibrated with the same buffer. After elution with a linear NaCl gradient (50-1000 mM in buffer LS), suitable fractions were pooled and dialyzed against buffer B6. Subsequently, the protein was concentrated, aliquoted, snap-frozen and stored at −80°C.

#### _DipMLytM_

His_6_-SUMO-DipM(459-609) was overproduced in *E. coli* AI041 and purified by Ni-NTA affinity purification as described for full-length DipM. After elution from the HisTrap HP column, fractions containing the protein in high concentrations were pooled and dialyzed against buffer CB (50 mM Tris, 150 mM NaCl, 10% v/v glycerol, adjusted to pH 8 with HCl). The solution was then supplemented with 1 mM DTT and His_6_-Ulp1 protease (Marblestone et al., 2006) at a 1:1000 molar ratio relative to His_6_-SUMO-DipM(459-609) and incubated for 2 h at 4°C. Subsequently, it was passed through a 0.22 µm filter to remove potential precipitates and applied onto a 5 ml HisTrap HP column (GE Healthcare, USA) equilibrated with buffer BZ3 as described above. Flow-through fractions containing DipM(459-609) at high purity were pooled and dialyzed against buffer B6. Finally, the protein was concentrated, aliquoted, snap-frozen and stored at −80°C.

For crystallization, DipM(459-609) was further purified by size exclusion chromatography after removal of the SUMO tag. To this end, flow-through fractions from the second Ni-NTA affinity purification step containing the protein at high purity were concentrated and then applied onto a HiLoad 16/60 Superdex 75 prep grade column (GE Healthcare, USA) equilibrated with a buffer containing 50 mM HEPES, 100 mM NaCl, adjusted to pH 7.2 with NaOH. After elution at a flow rate of 0.5 ml/min, peak fractions were concentrated, aliquoted, snap-frozen and stored at −80°C.

#### SdpA

To purify His_6_-SdpA(21-699), cells of *E. coli* AI033 were grown in LB medium at 37°C and then shifted to 18°C prior to the induction of protein overproduction by the addition of 1 mM IPTG (at an OD_600_ of 0.6). After incubation of the culture at 18°C for another 18 h, the cells were harvested, washed with buffer BZ3 and stored at −80°C. For further processing, the thawed cells were resuspended in buffer BZ3 (2 ml per 1 g of cell pellet) supplemented with DNase I (10 µg/ml) and PMSF (100 µg/ml). After three passages through a French press at 16,000 psi, cell debris was removed by centrifugation at 30 000 xg for 30 min. The supernatant was then applied onto a 5 ml HisTrap HP column (GE Healthcare, USA) equilibrated with buffer BZ3. The column was washed with 10 column volumes (CV) of buffer BZ3, and protein was eluted with a linear imidazole gradient (20-250 mM in buffer BZ3) at a flow rate of 1 ml/min. Fractions containing His_6_-SdpA at high purity were pooled, concentrated and applied onto a HiLoad 16/60 Superdex 200 prep grade size exclusion chromatography columns equilibrated with buffer B6. The peak fractions were concentrated, aliquoted, snap-frozen and stored at −80°C.

#### SdpB

His_6_-SUMO-SdpB(26-536) was overproduced in *E. coli* AI041 and purified by Ni-NTA chromatography as described for His_6_-SdpA(21-699). After elution from the HisTrap HP column, fractions containing the protein in high concentrations were pooled and dialyzed against buffer CB. The solution was supplemented with 1 mM DTT and His6-Ulp1 protease and incubated for 2 h at 4°C. Subsequently, it was passed through a 0.22 µm filter to remove potential precipitates and applied onto a 5 ml HisTrap HP column (GE Healthcare, USA) equilibrated with buffer BZ3. Flow-through fractions containing the protein at high purity were pooled and dialyzed against buffer. Finally, the protein was concentrated, aliquoted, snap-frozen and stored at −80°C.

#### AmiC

His_6_-SUMO-AmiC(35-395) was overproduced in *E. coli* AI061 and purified as described for His_6_-SUMO-SdpB(26-536).

#### _FtsNPeri_

His_6_-SUMO-FtsN(51-266) was overproduced in *E. coli* AI060 and isolated as described for His_6_-SUMO-DipM(459-609), with the addition of a size exclusion chromatography step to improve the purity of the preparation. After removal of the SUMO tag, fractions from the second Ni-NTA affinity purification step that were highly enriched in FtsN(51-266) were concentrated and applied to a HiLoad 16/60 Superdex 75 prep grade column (GE Healthcare, USA) equilibrated with buffer B6. Fractions containing the protein at high purity were pooled, concentrated, aliquoted, snap-frozen and stored at −80°C.

#### _CrbASPOR_

His_6_-SUMO-CrbA(371-451) was overproduced in *E. coli* AI075 and processed as described for His_6_-SUMO-DipM(459-609), with the addition of a last step in which the protein was dialyzed against buffer LS and then applied onto a HiTrap SP cation exchange column (GE Healthcare, USA) equilibrated in the same buffer. CrbA(371-451) was eluted with a linear NaCl gradient (50-1000 mM in buffer LS). Fractions containing the protein at high purity were pooled and dialyzed against buffer B6. Subsequently, they were concentrated, aliquoted, snap-frozen and stored at −80°C.

### Protein crystallization and structural analysis

High-throughput crystallization trials were performed in 96-well plates at 18°C in sitting drops consisting of 250 nL of protein solution (13.6 mg/mL) and 250 nL of precipitation solutions from commercial crystallization screens (Hampton Research, USA), using an Innovadine crystallization robot. Conditions producing crystals were optimized and scaled up to 1 µl of protein solution and 1 µl of precipitant against 150 µl of the crystallization solution in the reservoir. DipM crystallized in 0.02 M magnesium chloride, 0.1 M HEPES pH 7.5 and 22% polyacrylic acid sodium salt 5100. Crystals were cryoprotected in 30% polyethylene glycol prior to flash freezing. Diffraction data were collected at the ALBA synchrotron (XALOC beamline) using the Pilatus 6M detector. DipM crystals belonged to the P22_1_2_1_ space group with the dimensions *a*= 65.869, *b*= 105.839, *c*= 108.431, α= β=γ= 90°, and diffracted up to 2.25 Å resolution. The datasets were processed with XDS (Kabsch, 2010) and Aimless (Evans and Murshudov, 2013). The asymmetric unit was formed by four monomers, with 56.09% of solvent content. Structure determination was performed by molecular replacement, using Zoocin A from *Streptococcus equi* (PDB: 5KVP; 46.67% of sequence identity with DipM) as a search model. The model obtained was then completed manually using Coot (Emsley et al., 2010), followed by refinement using PHENIX (Liebschner et al., 2019) and REFMAC5 (Murshudov et al., 2011). A summary of the refinement statistics is given in **Supplementary Table 1**. Structures were visualized with ChimeraX (Goddard et al., 2018).

### Co-immunoprecipitation analysis

To identify interactors of DipM, SdpA, AmiC, LdpF and CrbA, we used strains producing FLAG-tagged bait proteins or untagged versions thereof (as negative controls) under the control of the xylose-inducible *P_xyl_* promoter in place of the respective native proteins. In the case of FtsN, a strain bearing *gfp* fused the endogenous *ftsN* gene was employed, and the wild-type strain NA1000 was used as negative control. The cells were grown in 500 ml (for DipM and SdpA) or 200 ml (for all remaining proteins) M2G medium, supplemented with 0.3% xylose when necessary, until they reached an OD_600_ of 0.6. After the addition of formaldehyde to a final concentration of 0.6%, the cultures were incubated for 5 min at room temperature, before the crosslinking reaction was stopped by the addition of glycine to a final concentration of 125 mM. Cells were harvested by centrifugation (12,000 ×g, 4°C, 10 min), washed with wash buffer (50 mM sodium phosphate pH 7.4, 5 mM MgCl_2_), pelleted and stored at −80°C. For further processing, the pellets were resuspended in 10 ml of a buffer containing 20 mM HEPES pH 7.4, 100 mM NaCl, 20 % (v/v) glycerol, 10 mg/ml lysozyme, 5 μg/ml DNase I, 100 μg/ml PMSF and 0.5% (for DipM and SdpA) or 1% (remaining proteins) of Triton X-100. The suspensions were then incubated for 30 min on ice and disrupted by three passages through a French press. The lysate was centrifuged (13,000 ×g, 4°C, 5 min) to remove intact cells and cell debris, and the supernatant was incubated with magnetic affinity beads carrying anti-FLAG (Sigma-Aldrich) or anti-GFP (Chromotek, Germany) antibodies for 2 h at 4°C in a rotator. The beads were then collected by centrifugation (4,000 ×g, 4°C) and resuspended in 700 µl of 100 mM ammonium-bicarbonate. After vigorous agitation, they were washed three times in 100 mM ammoniumbicarbonate using a magnetic separator. After removal of the supernatant of the last wash, the beads were resuspended in 100 µl of elution buffer 1 (1.6 M urea, 100 mM ammoniumbicarbonate, 5 µg/ml trypsin) and incubated for 30 min in a thermomixer (27°C, 1200 rpm). After collection of the beads in a magnetic separator, the supernatant was transferred to a new tube. The beads were resuspended in 40 µl of elution buffer 2 (1.6 M urea, 100 mM ammoniumbicarbonate, 1 mM tris[2-carboxyethyl]phosphine) and collected again. Subsequently, the supernatant was combined with the previous eluate, and the elution with elution buffer 2 was repeated one more time. The pooled fractions were left overnight at room temperature. On the following day, 40 µl of iodoacetamide (5 mg/ml) were added, and the mixture was incubated for 30 min in the dark. After the addition of 150 µl of 5% [v/v] trifluoroacetic acid (TFA), the mix was passed through C-18 microspin columns (Harvard Apparatus), previously conditioned with acetonitrile and equilibrated with buffer A (0.1% [v/v] TFA in water). The column was then washed three times with 150 µl of buffer C (5% [v/v] acetonitrile, 95% [v/v] water and 0.1% [v/v] TFA). The peptides were eluted by washing the column three times with 100 µl of buffer B (50% [v/v] acetonitrile, 49.9% water [v/v] and 0.1% [v/v] TFA). The combined eluates were then dried under vacuum, and peptides were suspended in LC buffer (0.15% [v/v] formic acid, 2% [v/v] acetonitrile) by 20 pulses of ultrasonication (Amplitute 20, Cycle 0.5) and shaking for 5 min at 1,400 rpm and 25°C.

LC-MS analysis of the peptide samples was carried out on a Q-Exactive Plus instrument connected to an Ultimate 3000 RSLC nano and a nanospray flex ion source (all Thermo Scientific). Peptide separation was performed on a reverse phase HPLC column (75 μm x 42 cm) packed in-house with C18 resin (2.4 μm, Dr. Maisch). The peptides were loaded onto a PepMap 100 precolumn (Thermo Scientific) and the eluted by a linear acetonitrile gradient (2-35% solvent B) over 60 or 90 min (solvent A: 0.15% [v/v] formic acid in water, solvent B: 0.15% formic acid [v/v] in acetonitrile). The flow rate was set to 300 nl/min. The spray voltage was set to 2.5 kV, and the temperature of the heated capillary was set to 300°C. Survey full-scan MS spectra (m/z = 375-1500) were acquired in the Orbitrap with a resolution of 70,000 full width at half maximum at a theoretical m/z 200 after accumulation of 3×106 ions in the Orbitrap. Based on the survey scan, up to 10 of the most intense ions were subjected to fragmentation using high collision dissociation (HCD) at 27% normalized collision energy. Fragment spectra were acquired at 17,500 resolution. The ion accumulation time was set to 50 ms for both MS survey and MS/MS scans. To increase the efficiency of MS/MS attempts, the charged state screening modus was enabled to exclude unassigned and singly charged ions. The dynamic exclusion duration was set to 30 sec.

The resulting raw data were analyzed using Mascot (v 2.5, Matrix Science). Search results were loaded into Scaffold 4 (Proteome Software) to extract total spectrum counts for further analysis. The peptide count data were loaded in Perseus (Tyanova et al., 2016) (version 1.5.8.5) to generate volcano plots. In brief, one unit was added to all the counts to eliminate the zeroes and then log2 was applied to all data. Columns were classified according to whether they belonged to the sample or negative control, and volcano plots were generated using the default settings. The resulting data on enrichment (difference to negative control) and significance (-log10 of *p* value) were exported to Microsoft Excel 2019, where they were re-plotted to generate the figures.

### Biolayer interferometry

Bio-layer interferometry experiments were conducted in a BLItz system (PALL Life sciences, USA) equipped with High Precision Streptavidin (SAX) biosensors (PALL Life sciences, USA). Proteins were biotinylated using NHS-PEG4-Biotin (Thermo Scientific, USA) following the manufacturer recommendations. After immobilization on the sensor surface and the establishment of a stable baseline, the biotinylated proteins were probed with the indicated analytes. After the binding step, the sensors were transferred to an analyte-free buffer to follow the dissociation kinetics. The extent of non-specific interactions was determined by analyzing the binding of the respective analyte to unmodified sensors. All BLI analyses were performed in a buffer containing 50 mM HEPES/NaOH pH 7.2, 50 mM NaCl, 5 mM MgCl_2_, 0.1 mM EDTA, 10% glycerol, 10 µM BSA, 0.01% Triton X-100.

For analysis, BLI data were processed in Microsoft Excel 2019 using the Solver add-in. In brief, the association/dissociation curves were normalized against the baselines obtained prior to exposure of the sensor to the analyte. The wavelength shift values reached after equilibration of the binding reactions were fitted to a one-site binding model to calculate the equilibrium dissociation constants (*K*_D_).

### Peptidoglycan digestion assays

To assay for lytic transglycosylase activity, SdpA (5 μM) alone or in combination with DipM or DipM^LytM^ (5 μM) was incubated with purified peptidoglycan (0.5 mg/ml) from *E. coli* MC1061 (Casadaban and Cohen, 1980) in 20 mM HEPES/NaOH pH 7.5, 20 mM NaCl, 1 mM MgCl_2_ in a total volume of 50 μl for 4 h in a thermal shaker set to at 37°C and 900 rpm. In parallel, a control reaction containing no SdpA was performed. The reactions were stopped by boiling the samples for 10 min at 100°C in a dry-bed heat block. The samples were centrifuged at room temperature for 15 min at 16,000 ×g. The supernatant was recovered and the pH adjusted to 3.5-4.5 with 20% phosphoric acid. The samples were analysed by HPLC as published previously (Glauner, 1988) but with a modified buffer B and a linear gradient over 140 min from buffer A (50 mM sodium phosphate pH 4.31 with 1 mg/L sodium azide) to buffer B (75 mM sodium phosphate pH 4.95, 30% methanol). Eluted muropeptides were detected by their absorbance at 205 nm. An overnight digestion of peptidoglycan with Slt (5 µM) (Banzhaf et al., 2020) from *E. coli* MC1061 served as a positive control. Peaks were assigned by their retention time and quantified by UV absorbance. Statistical tests comparing the integrated peak areas, were performed in Microsoft Excel 2019.

To assay for amidase activity, peptidoglycan (0.5 mg/ml) from *E. coli* MC1061 was incubated with AmiC alone and with AmiC combined with either DipM or DipM^LytM^ (all 5 μM) in 20 mM Hepes/NaOH pH 7.5, 20 mM NaCl in a total volume of 50 μl for 16 h in a thermal shaker set to 37°C and 900 rpm. Control reactions did not contain AmiC. The samples were boiled at 100°C for 10 min in a dry-bed heat block, acidified to pH 4.8 and incubated further with 10 μg of cellosyl (Hoechst, Frankfurt am Main, Germany) for 16 h at 37°C in a thermal shaker. Subsequently, the samples were boiled for 10 min in a dry-bed heat block and centrifuged at room temperature for 15 min at 16,000 ×g. After retrieval of the supernatant, the muropeptides in the supernatant were reduced with sodium borohydride and separated by HPLC analysis as described previously (Glauner, 1988).

## Acknowledgements

We thank Aleksandra Zielińska for help in the initial phases of the project, Manuel Osorio-Valeriano for advice on protein purification and biochemistry, Julia Rosum for excellent technical assistance, Daniela Vollmer for the purification of peptidoglycan, and the staff of the ALBA Synchrotron facilities for support during crystallographic data collection. This work was supported by the University of Marburg (core funding to P.G. and M.T.), the Max Planck Society (Max Planck Fellowship to M.T.), the German Research Foundation (DFG; project 269423233 – TRR 174 to P.L.G.), the United Kingdom Research and Innovation (UKRI) Strategic Priorities Fund (grant EP/T002778/1 to W.V.) and the Spanish Agency of Research at the Ministry of Science and Innovation (grant PID2020-115331GB-I00 to J.A.H.). A.I.-M. was a fellow of the International Max Planck Research School for Environmental, Cellular and Molecular Microbiology (IMPRS-Mic).

## Data availability

The atomic coordinates for the crystal structure of DipM^LytM^ were deposited in the RCSB Protein Data Bank with the accession number 7QRL. All other data generated in this study are included in the manuscript and the supplemental information.

## Author contributions

A.I-M. and M.T. conceived the study. A.I-M. constructed the plasmids and strains, purified the proteins, performed the Co-IP and BLI-based interaction analyses and conducted the growth analyses and subcellular localization studies. V.M.-R. and M.T.B. performed the crystallization screens, solved the crystal structure of DipM^LytM^ and performed the molecular modeling studies. R.H.-T. conducted the single-particule tracking analyses. J.B. performed the peptidoglycan digestion assays. T.G. performed the mass spectrometric analyses. A.I.-M., V.M.-R., R.H.-T., J.B., M.T.B., T.G., W.V., J.A.H and M.T. analyzed the data. M.B., W.V., P.L.G., J.A.H. and M.T. supervised the study. W.V., P.L.G., J.A.H. and M.T. secured funding. A.I.-M. and M.T. wrote the paper, with input from all other authors.

## Supplementary material

### Supplementary figures

**Figure S1.**
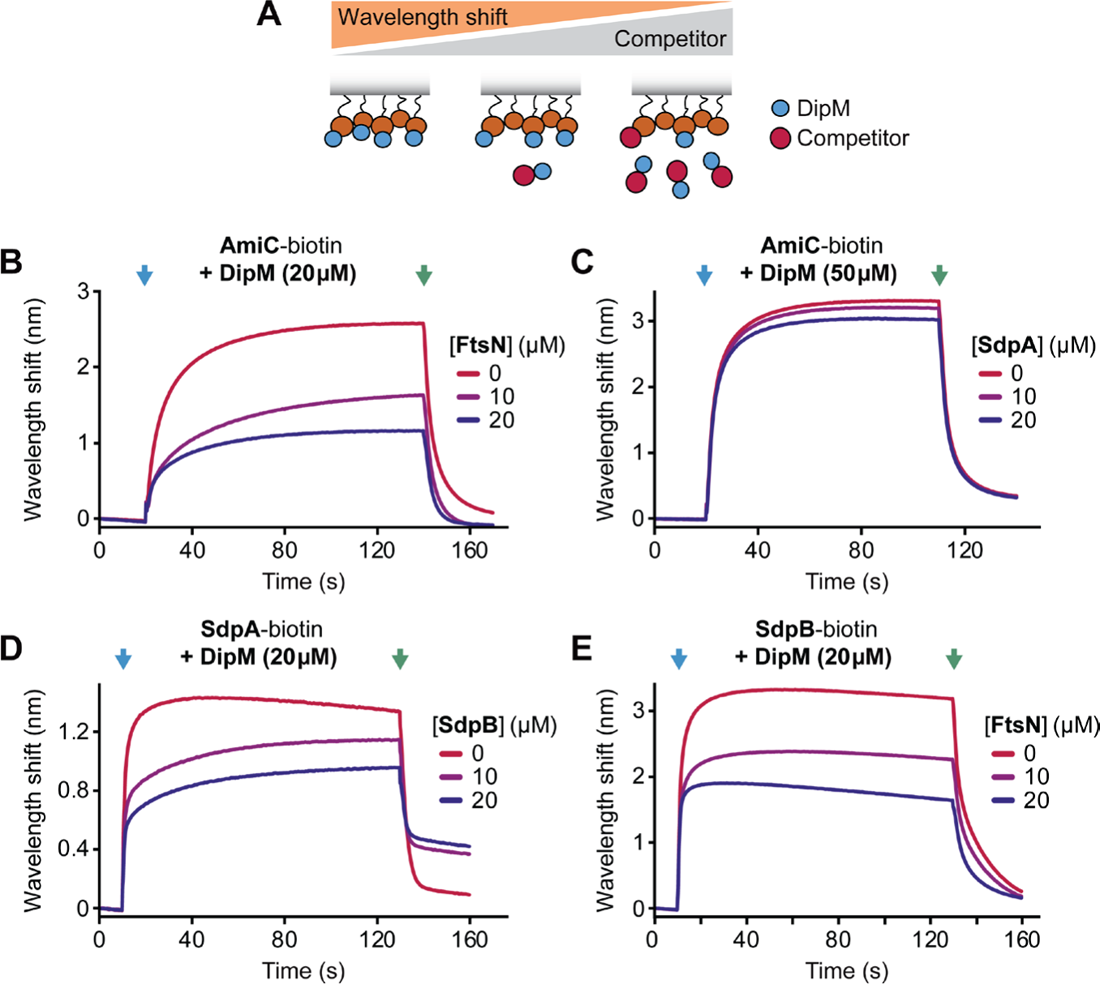
The different interactors compete for binding to DipM. **(A)** Schematic representation of the BLI-based competition assay used in this study. After immobilization of one of the interactors on the sensor surface, the sensor is probed with DipM alone or with mixtures of DipM and a second interactor. If the two interactors bind to different, non-overlapping sites on DipM, they form a ternary complex on the sensor, leading to an increase in the wavelength shift. Otherwise, the signal remains largely constant or decreases with increasing concentrations of the second interactor, depending on the relative affinities of the interactions. **(B-E)** Competitive binding of two interactors to DipM. Sensors derivatized with the indicated biotinylated interactor were probed with DipM alone or with a mixture of DipM preincubated with a second interactor at the indicated concentrations. A blue arrow marks the start of the association phase, a green arrow the start of the dissociation phase. All assays were performed at least in duplicate, with similar results obtained throughout.

**Figure S2.**
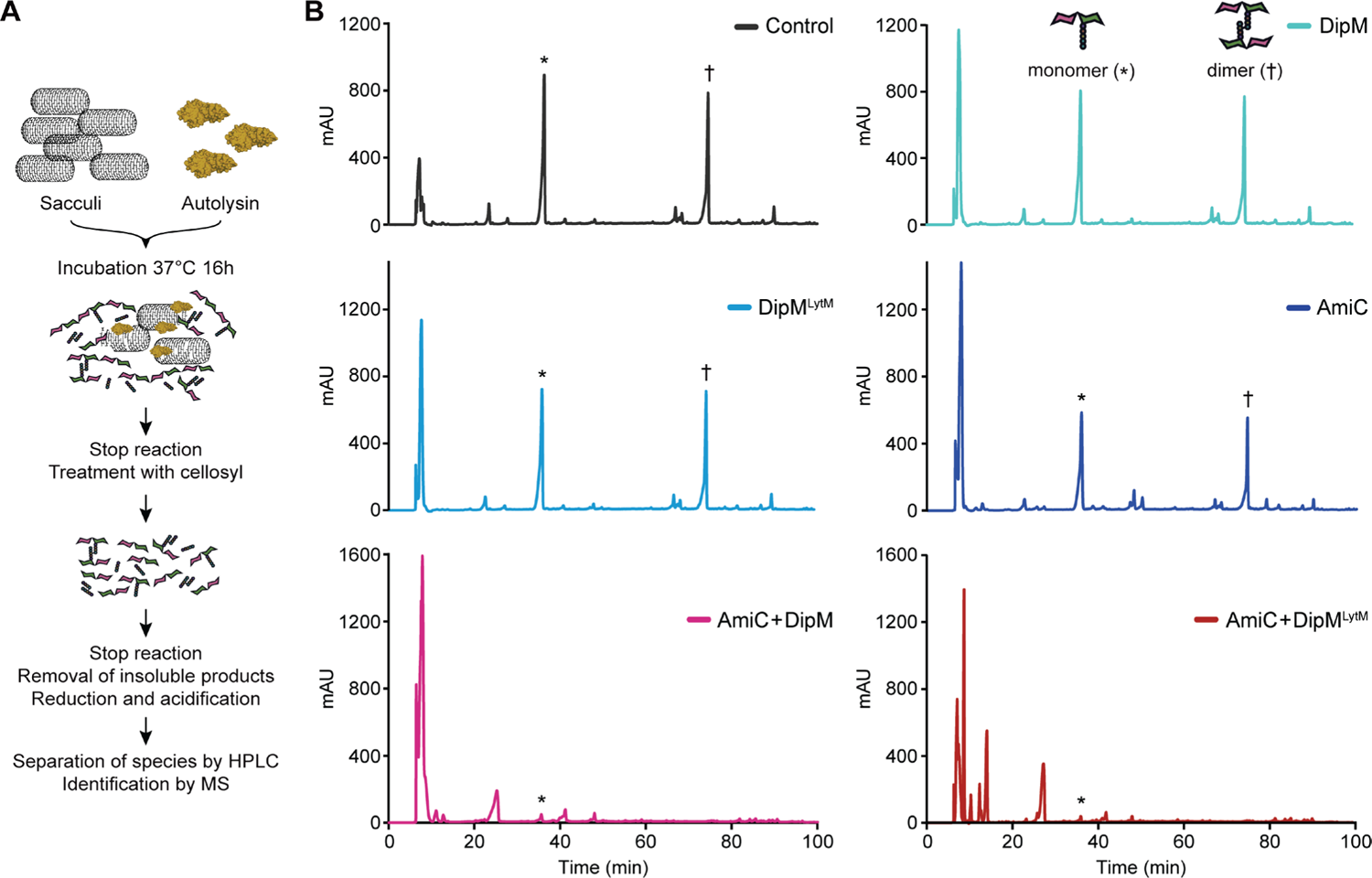
DipM strongly stimulates the amidase activity of AmiC. **(A)** Overview of the procedure used to assess the activity of AmiC. **(B)** HPLC chromatograms showing the muropeptides generated by the treatment of sacculi with cellosyl (Control) and changes in the muro-peptide profile resulting from the subsequent incubation of these muropeptides with the indicated protein(s). AmiC and DipM/DipM^LytM^ were used at equimolar ratios. Asterisks mark the peaks of monomeric products, daggers those of dimeric products. Amidase activity is indicated by a decrease in the abundance of these species, because the reaction generates free sugar dimers and peptides, which do not bind to the column under the conditions used.

**Figure S3.**
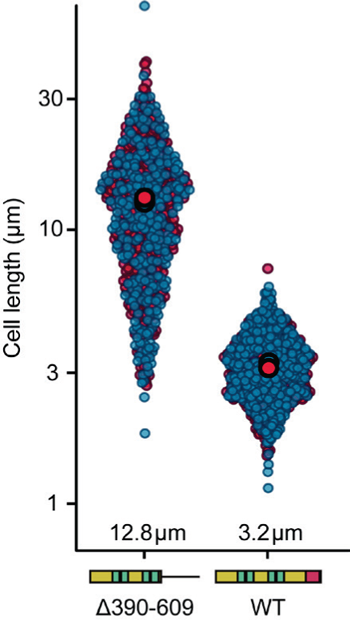
A truncated DipM variant lacking the LytM domain fails to complement the lack of native DipM. Superplots representing the cell length distribution in populations of cells producing DipM-sfmTurquoise2^ox^ (AI093) or a derivative lacking the LytM domain (Δ390-609; AI103) in place of the native protein in 2xPYE medium. The data represent the results of three replicates, which are displayed in different colors (red and blue). The big filled circles indicate the mean cell length obtained for each replicate.

**Figure S4.**
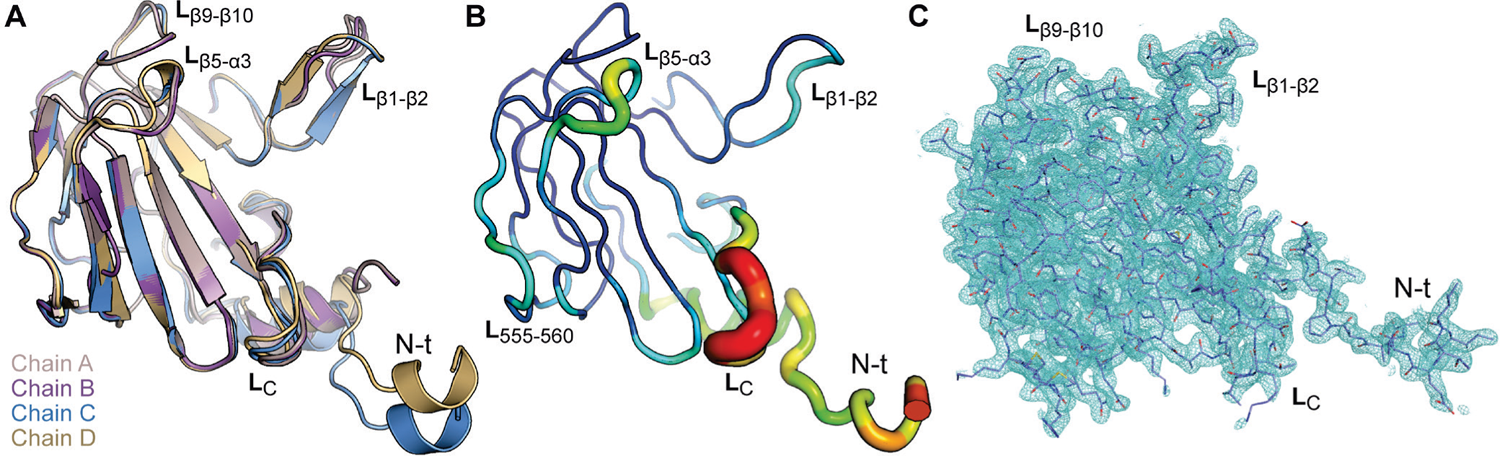
Electron density, structural plasticity and B-factor distribution for the DipM^LytM^ monomer. **(A)** Superimposition of the four independent molecules (chains A-D) constituting the asymmetric unit. Monomers are shown in cartoon view and colored differently. Variable regions are labeled. **(B)** Putty tube representation of the B-factor for the DipM^LytM^ reference chain (chain C). The color of the backbone varies depending on the B-factor of the residues, ranging from blue (lowest) to red (highest). In addition, the diameter of the tube increases with the size of the B-factor. **(C)** 2Fo-Fc electron density map for DipM^LytM^ chain C contoured at 1σ (shown as a blue mesh). Relevant regions are labeled.

**Figure S5.**
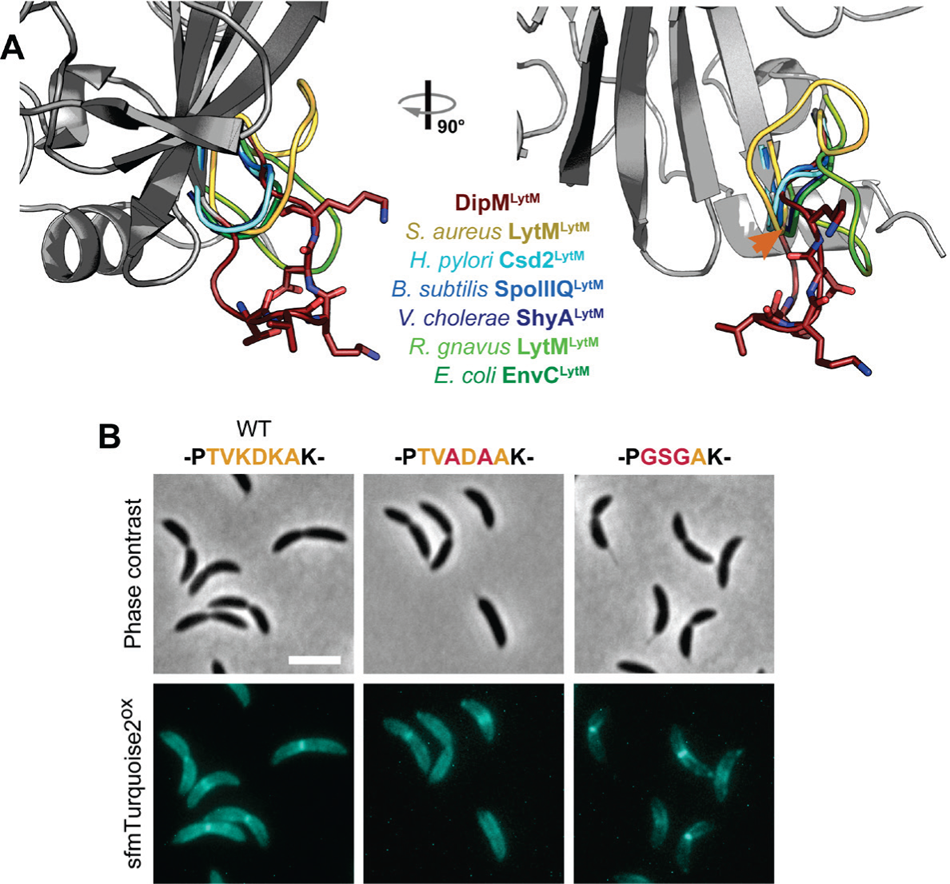
Role of the DipM-specific loop at the C-terminal end of DipM^LytM^. **(A)** Structural alignment of the crystal structures of DipM^LytM^ and the LytM domains of six other proteins, including *S. aureus* LytM (PDB: 4ZYB) (Grabowska et al., 2015), *H. pylori* Csd2 (PDB: 5J1L) (An et al., 2016), *B. subtilis* SpoIIIQ (PDB: 3UZ0) (Meisner et al., 2012), *V. cholerae* ShyA (PDB: 6UE4) (Shin et al., 2020), *R. gnavus* LytM (PDB: 3NYY) and *E. coli* EnvC (PDB: 4EH5). For all proteins except for DipM, only the loop following the last β-sheet of each LytM domain is shown for clarity, represented as colored ribbons without any secondary structural elements. The residues in the KDK motif, which is conserved in *C. crescentus* and close relatives, are shown in stick representation. The orange arrowhead indicates the position at which, in most structures, the loop turns upwards. **(B)** Functionality of DipM variants with exchanges in the conserved KDK motif. Shown are phase contrast and fluorescence images of cells producing the indicated DipM-sfmTurquoise2^ox^ variants in place of the native protein in 2xPYE medium (AI111, AI115). The native sequence of the conserved loop (orange) and residues exchanged in the mutant variants (red) are given on top of the corresponding images.

**Figure S6.**
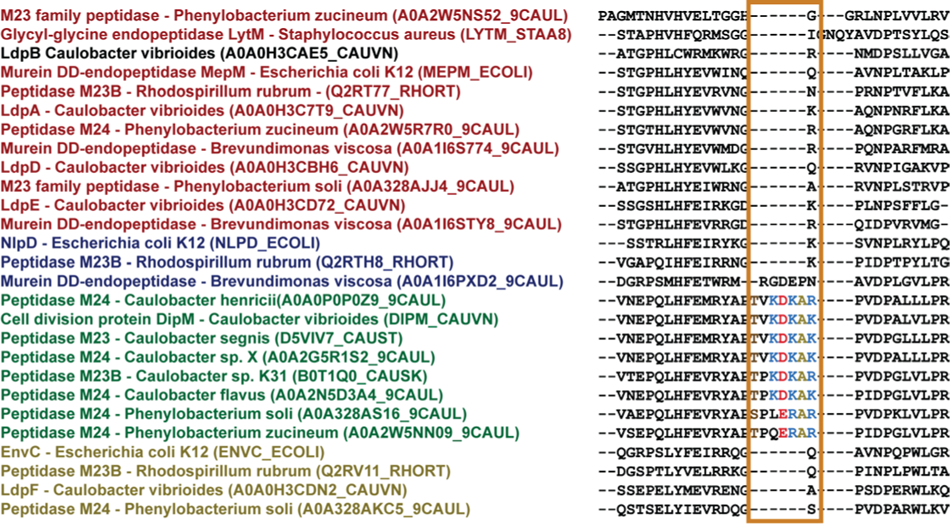
Alignment of the C-terminal regions of DipM and other LytM domain-containing proteins. DipM homologs of the genera *Caulobacter* and *Phenylobacterium* are shown in green, NlpD homologs of alpha- and gammaproteobacteria in blue, EnvC homologs of alpha- and gammaproteobacteria in olive, and catalytically active LytM domain-containing proteins from alpha- and gammaproteobacteria as well as LytM of *S. aureus* in red. *C. crescentus* LdpB, whose catalytic proficiency is still unclear, is shown in black. The conserved loop present in the *Caulobacter* and *Phenylobacterium* homologs is highlighted with an orange box and its conserved residues are shown in color.

**Figure S7.**
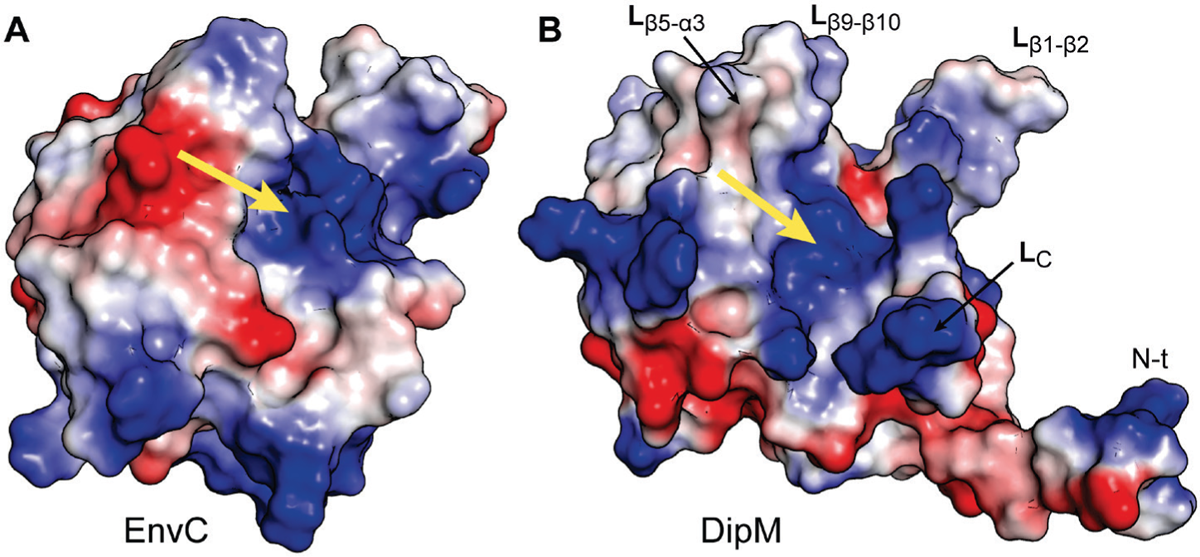
Positive electrostatic surface potential of the putative AmiC binding groove of DipM^LytM^. Shown is the electrostatic surface potential of **(A)** the LytM domain of EnvC and **(B)** DipM^LytM^, with positive and negative charges colored in blue and red, respectively. Yellow arrows point to the positive electrostatic potential in the binding groove. Loops delimiting the cavity in DipM are labeled.

**Figure S8.**
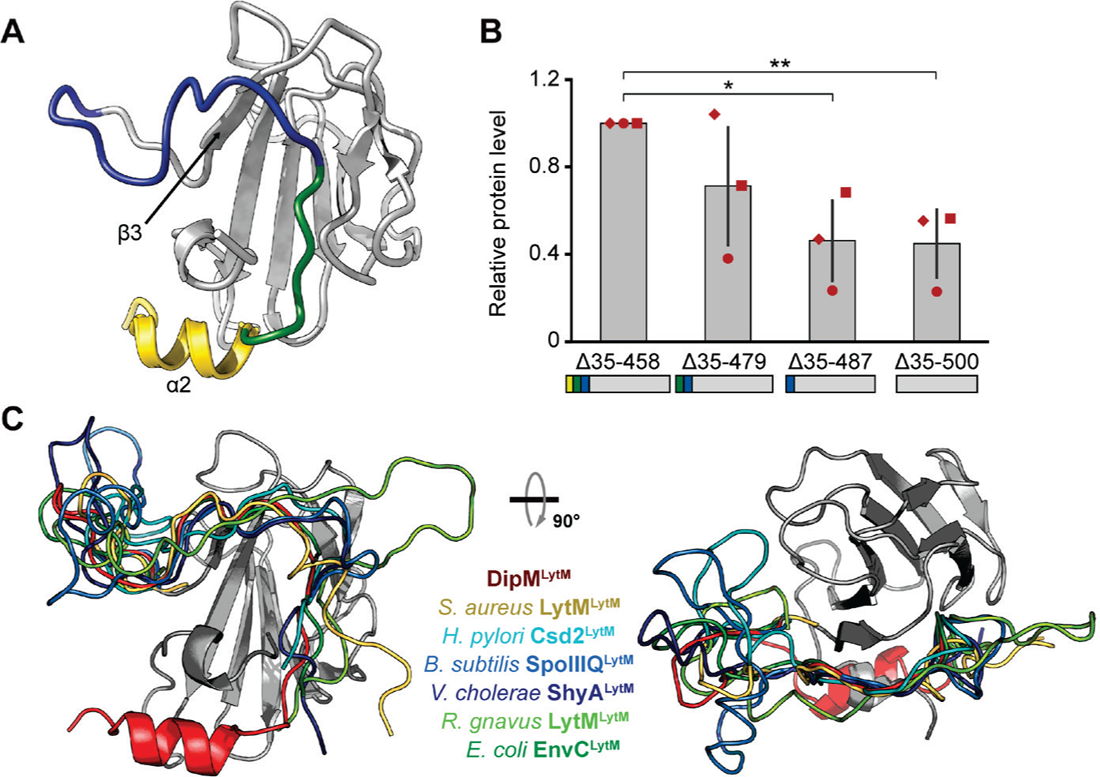
The N-terminal region of DipM^LytM^ is required for protein stability. **(A)** Shown is a cartoon representation of DipM^LytM^. The part of the protein that is recognized by the Hidden Markov Model employed to identify the LytM domain by the Pfam database (Mistry et al., 2021) is shown in grey. The remaining N-terminal region is divided into three parts: residues that are closer to the LytM domain and contact strand β2 and adjacent regions (blue), the following segment up to helix α2 (green) and helix α2 (yellow). **(B)** Bar chart representing the average (± SD) levels of the indicated DipM-sfmTurquoise2^ox^ variants (n=3), as determined by Western blot analysis of cells producing the fusion proteins in the wild-type background (AI098, AI123, AI124, AI125). The individual data points from the three replicates are shown as red symbols. The schematics at the bottom of the chart depict the architecture of the different protein variants. Asterisks indicate the statistical significance of differences between the averages obtained, determined by a one-way ANOVA test: **\*** p<0.05 and **\*\*** p<0.01. **(C)** Structural alignment of the crystal structures of DipM^LytM^ (in grey) and the LytM domains of six other proteins: *S. aureus* LytM (PDB: 4ZYB) (Grabowska et al., 2015), *H. pylori* Csd2 (PDB: 5J1L) (An et al., 2016), *B. subtilis* SpoIIIQ (PDB: 3UZ0) (Meisner et al., 2012), *V. cholerae* ShyA (PDB: 6UE4) (Shin et al., 2020), *R. gnavus* LytM (PDB: 3NYY) and *E. coli* EnvC (PDB: 4EH5). For all proteins except for DipM, only the N-terminal region adjacent to the LytM domain is shown for clarity, represented as colored ribbons without any secondary structural elements.

**Figure S9.**
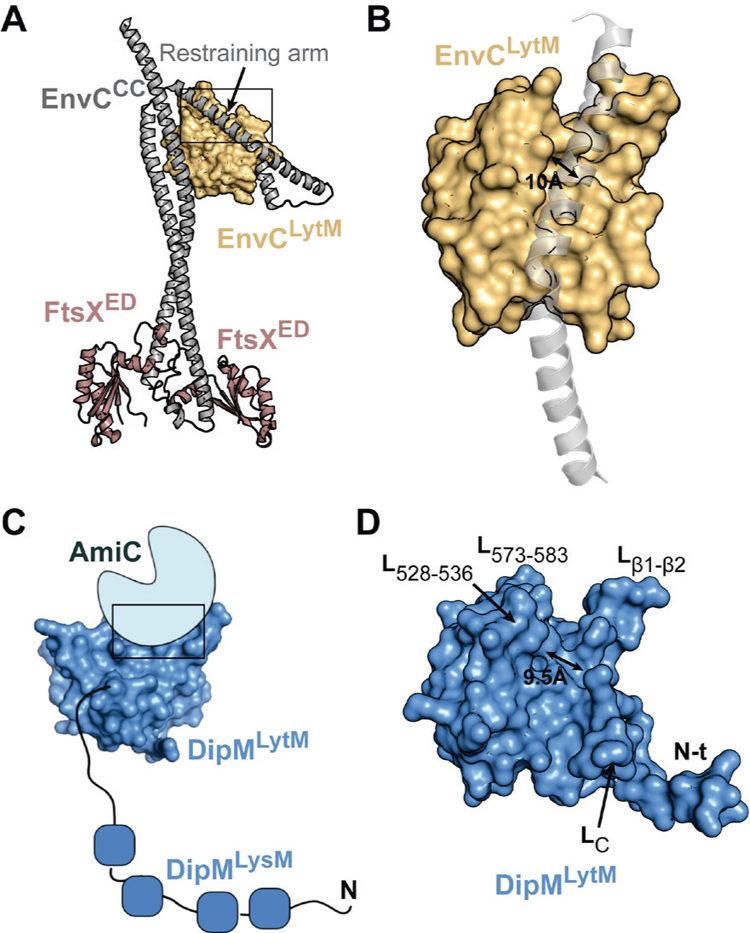
Comparison of the structures of EnvC and DipM. **(A)** Crystal structure of EnvC bound to the periplasmic domain of FtsX (FtsX^ED^). The LytM domain of EnvC (EnvC^LytM^) and its N-terminal coiled-coil region (EnvC^CC^) are indicated (PDB: 6TPI) (Cook et al., 2020). **(B)** Detailed view of the self-inhibitory structure form through interaction of EnvC^LytM^ (yellow) with the restraining arm (transparent gray cartoon), highlighted by a black box in panel A. **(C)** Schematic model of the DipM^LytM^-AmiC complex of *C. crescentus*, as predicted by AlphaFold-Multimer (Evans et al., 2022) (detailed in Figure 7C). **(D)** Surface view of DipM^LytM^, arranged in the same orientation as EnvC in panel B.

**Figure S10.**
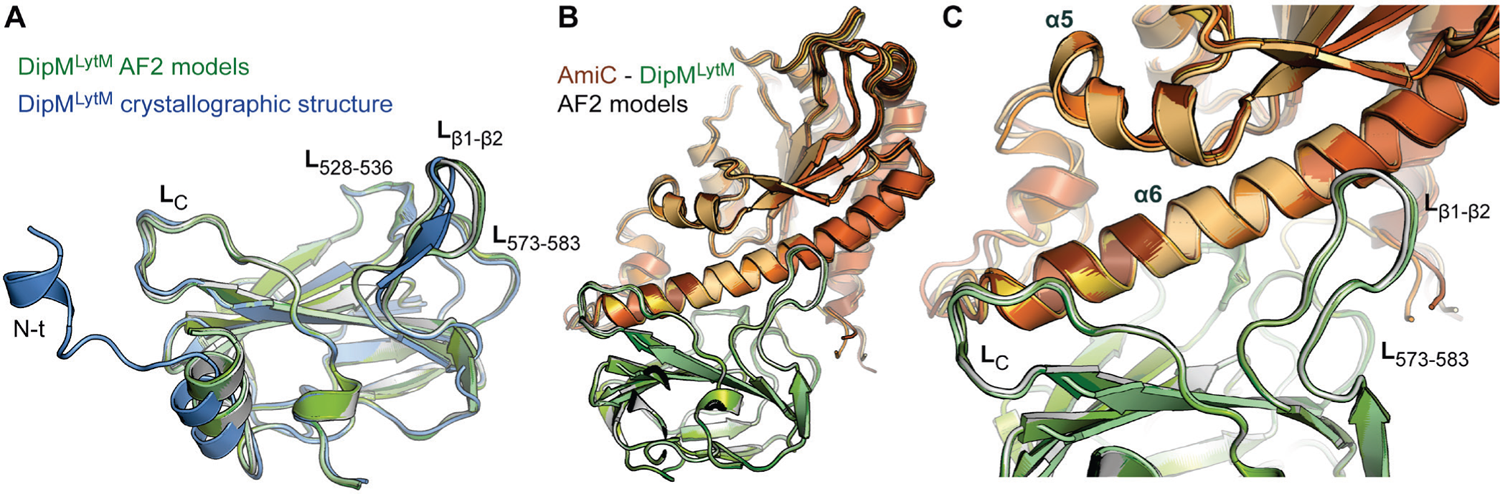
Evaluation of the model of the DipM^LytM^-AmiC complex generated by AlphaFold-Multimer. **(A)** Structural superimposition of the crystal structure of DipM^LytM^ (chain C, in blue) and a model of DipM^LytM^ generated by AlphaFold-Multimer (Evans et al., 2022) (in various shades of green). **(B)** Superimposition of DipM^LytM^-AmiC complexes predicted by AlphaFold-Multimer. DipM^LytM^ is shown in green, the different AmiC models in various shades of orange. **(C)** Magnified view of the predicted interacting regions of Dip^LytM^ and AmiC.

**Figure S11.**
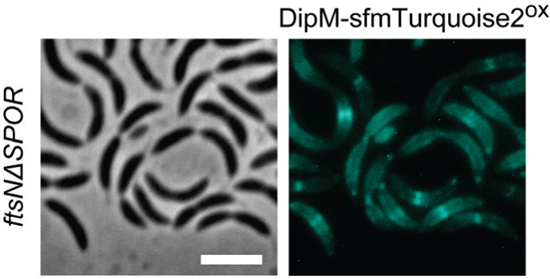
The SPOR domain of FtsN is not required for the recruitment of DipM to the division plane. Shown are representative phase contrast and fluorescence micrographs of cells producing DipM-sfmTurquoise2^ox^ in an *ftsNΔSPOR* background (AI117).

**Figure S12.**
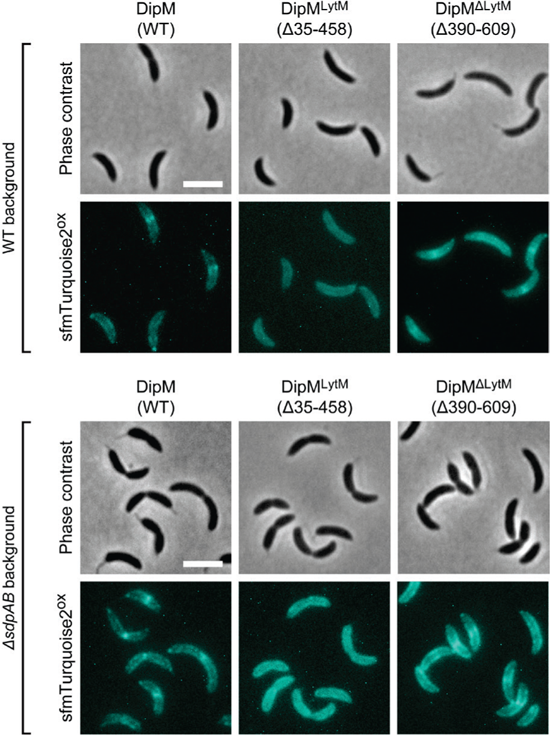
DipM localization is independent of the presence of SdpA/B. Phase contrast and fluorescence images of the strains used for single-molecule tracking (see Figure 9). Shown are representative phase contrast and fluorescence images of cells producing the indicated DipM-sfmTurquoise2^ox^ variants in the wild-type or Δ*sdpAB* backgrounds. The production of the fluorescent protein fusions was induced for 2 h with 0.3% xylose prior to microscopic analysis.

**Figure S13.**
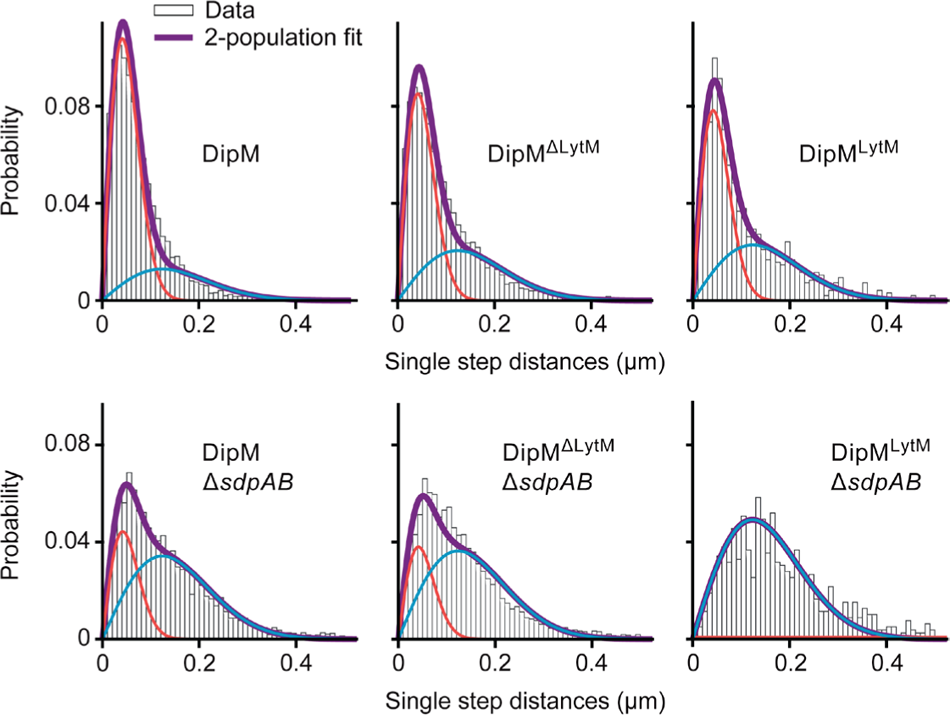
Single-molecule mobilities of different DipM variants. Shown is a Gaussian-mixture-analysis of the mobility of the indicated DipM-sfmTurquoise2^ox^ variants (measured by single-particle tracking as described in the legend to Figure 9). The probability distributions of the single-step frame-to-frame displacements obtained in the single-particle tracking experiments were fitted to a two-component Gaussian function, assuming a slow-moving (red line) and fast-moving (blue line) population.

### Supplementary tables

**Table S1.**
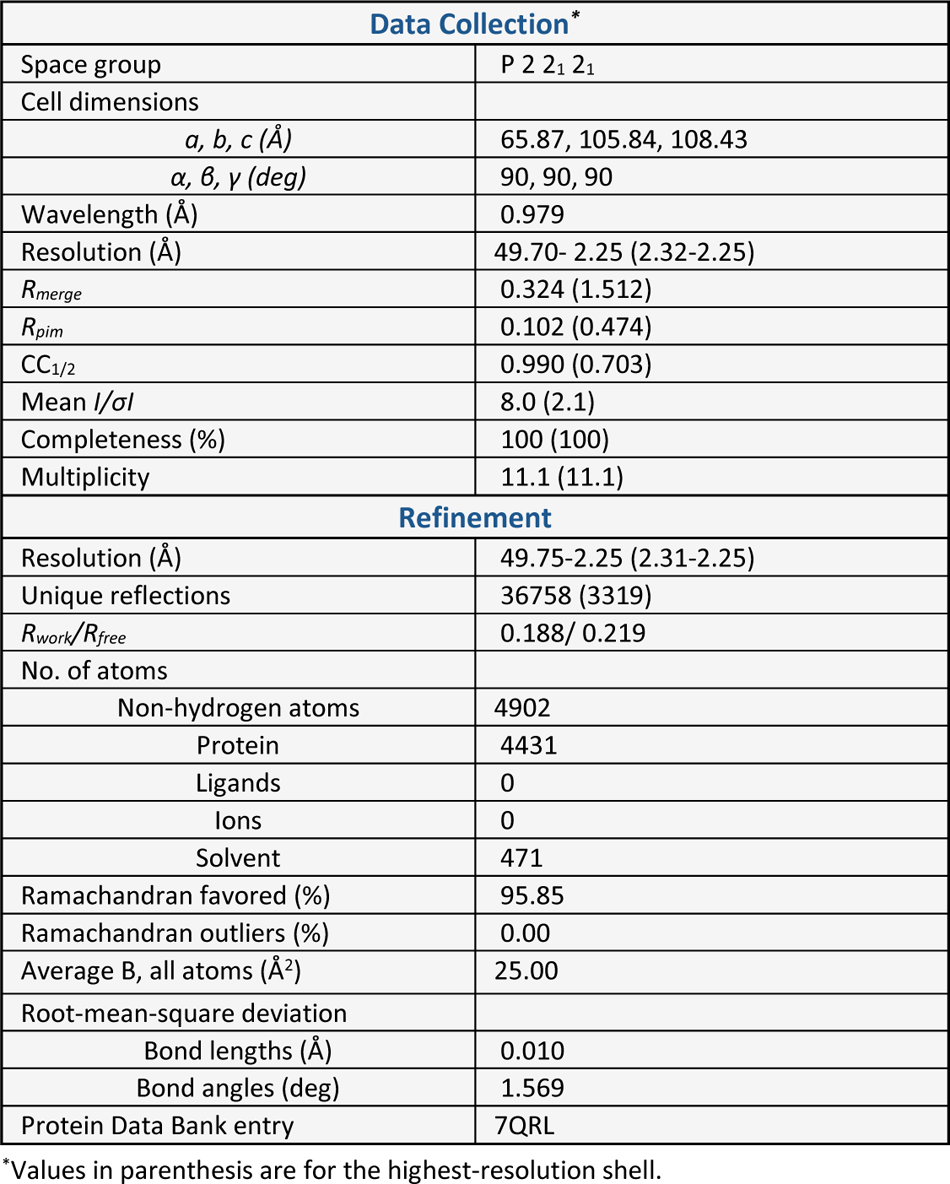
Data collection and refinement statistics for DipM^LytM^.

**Table S2.**
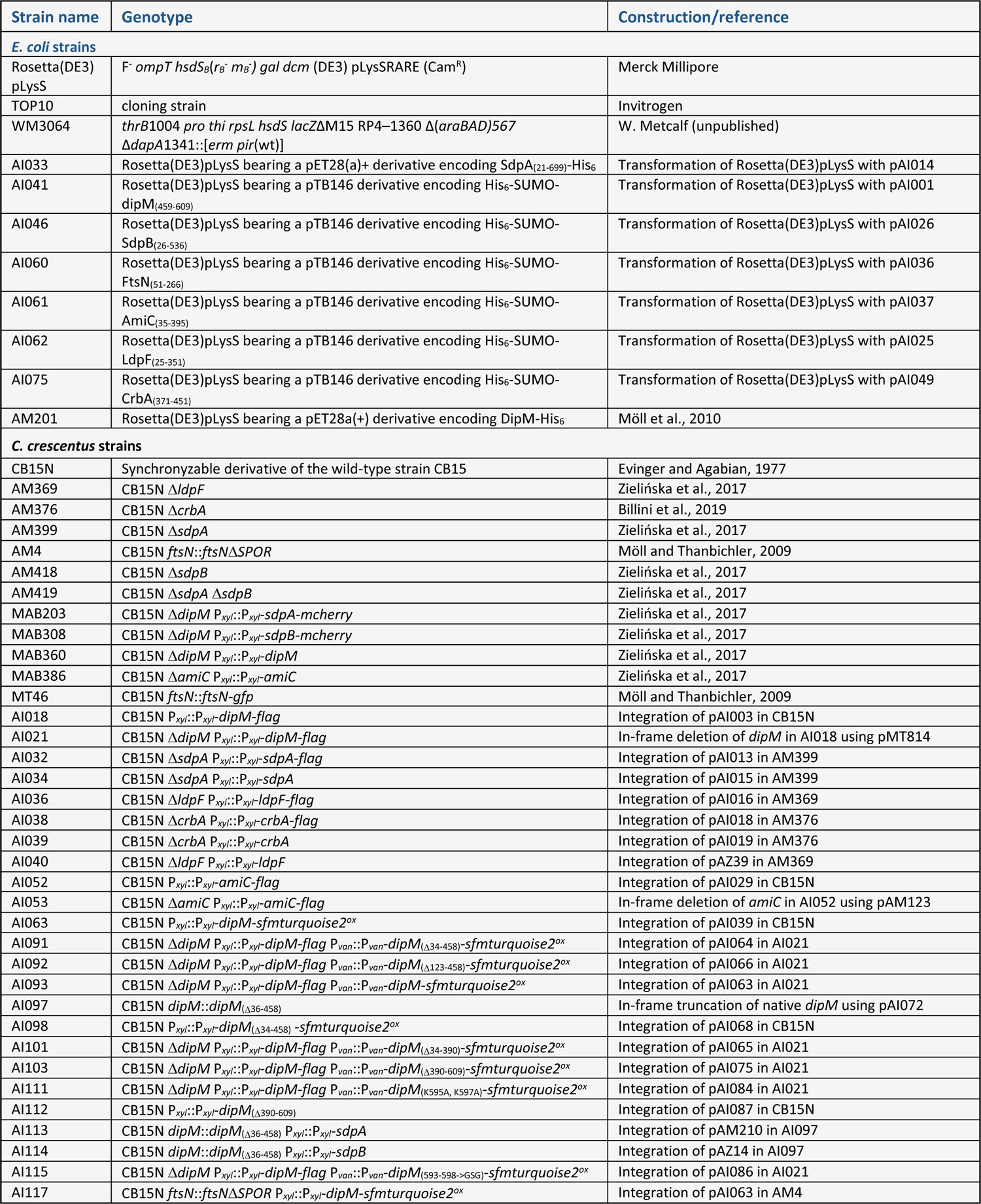
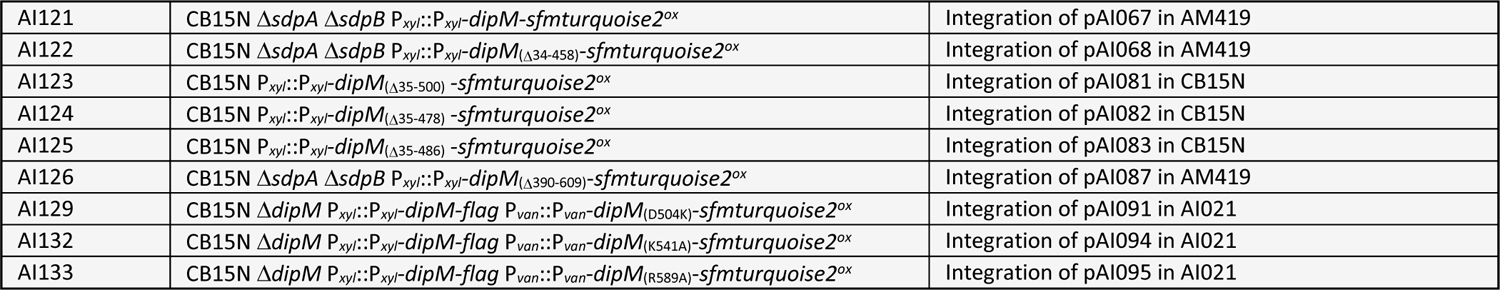
Strains used in this study.

**Table S3.**
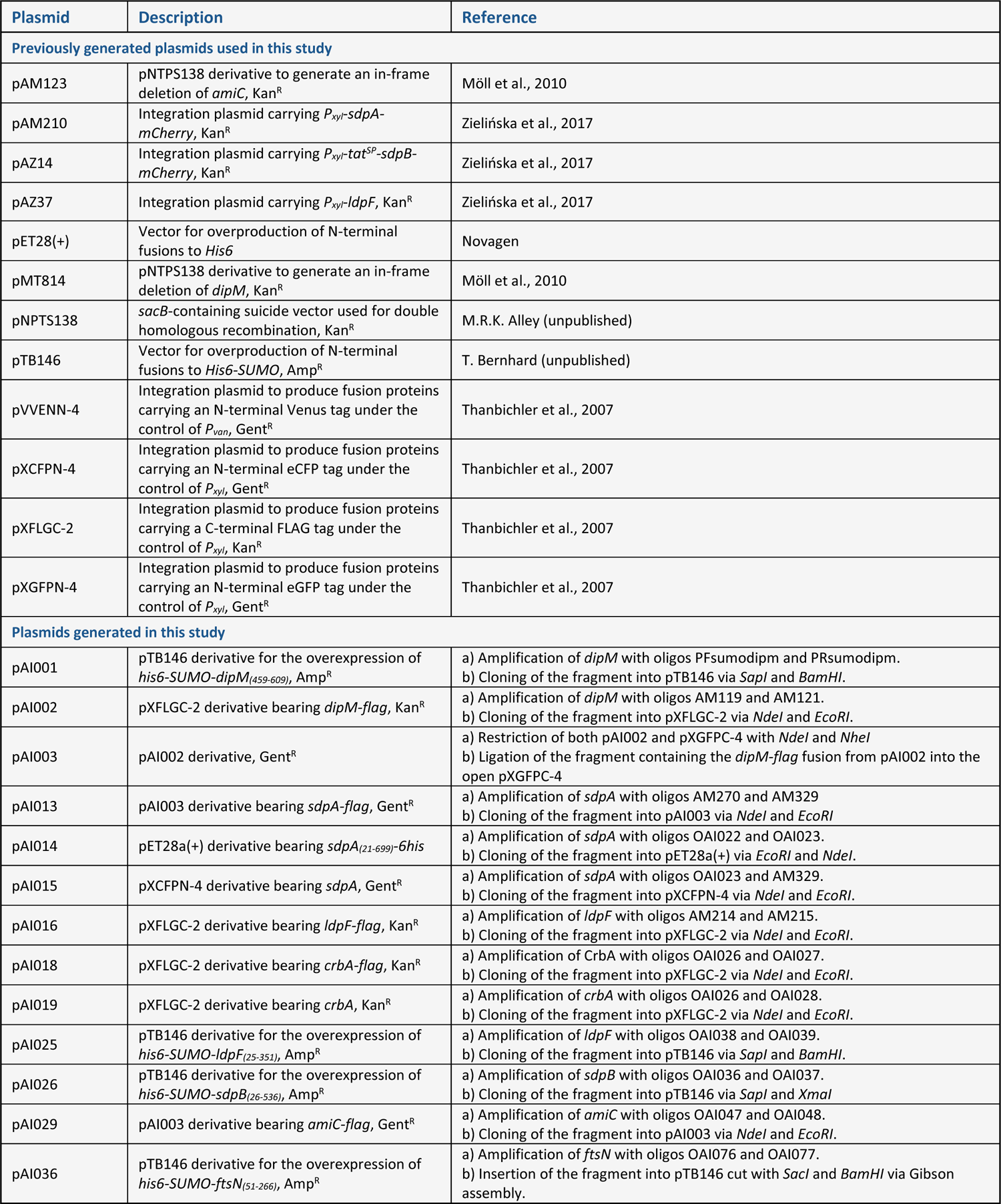
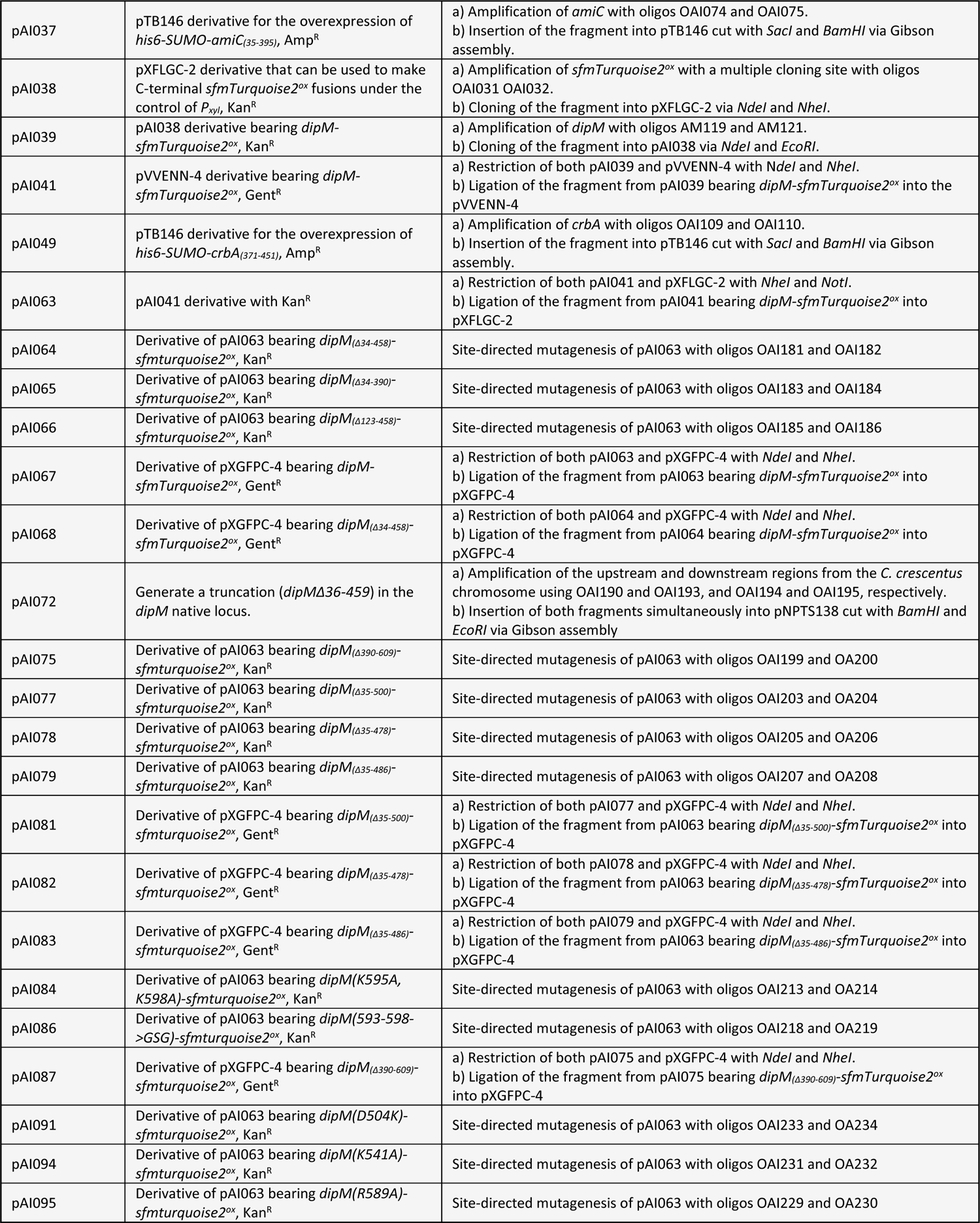
Plasmids used in this study.

**Table S4.**
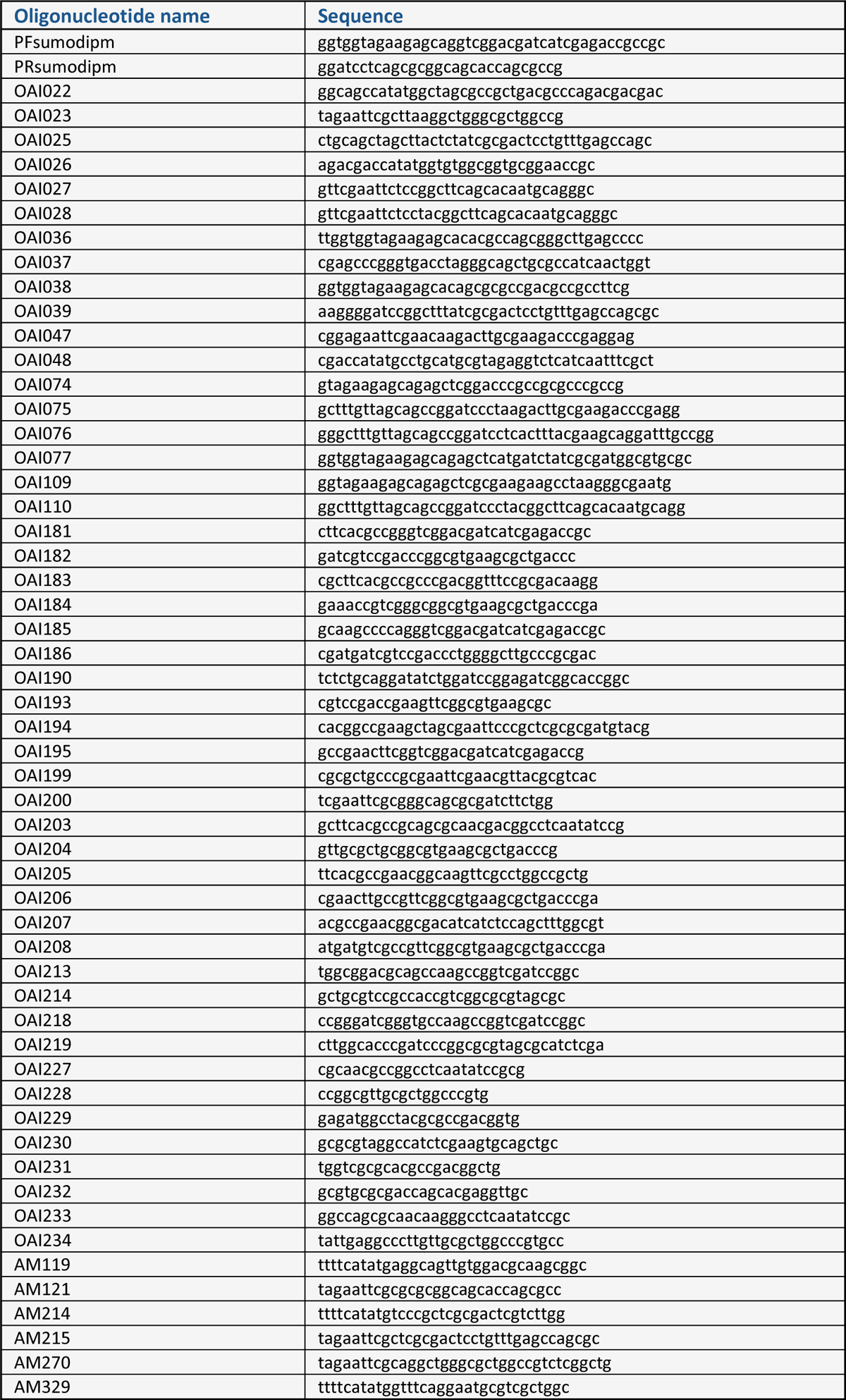
Oligonucleotides used in this work.

## References

1. Aaron, M, Charbon, G, Lam, H, Schwarz, H, Vollmer, W, Jacobs-Wagner, C (2007). The tubulin homologue FtsZ contributes to cell elongation by guiding cell wall precursor synthesis in *Caulobacter crescentus*. Mol. Microbiol. 64:938–952.

2. Alcorlo, M, Dik, DA, De Benedetti, S, Mahasenan, KV, Lee, M, Domínguez-Gil, T, Hesek, D, Lastochkin, E, López, D, Boggess, B, et al. (2019). Structural basis of denuded glycan recognition by SPOR domains in bacterial cell division. Nat. Commun. 10:5567.

3. Arends, SJR, Kustusch, RJ, Weiss, DS (2009). ATP-binding site lesions in FtsE impair cell division. J. Bacteriol. 191:3772–3784.

4. Banzhaf, M, Yau, HC, Verheul, J, Lodge, A, Kritikos, G, Mateus, A, Cordier, B, Hov, AK, Stein, F, Wartel, M, et al. (2020). Outer membrane lipoprotein NlpI scaffolds peptidoglycan hydrolases within multi-enzyme complexes in *Escherichia coli*. EMBO J. 39:e102246.

5. Bartual, SG, Straume, D, Stamsås, GA, Muñoz, IG, Alfonso, C, Martínez-Ripoll, M, Håvarstein, LS, Hermoso, JA (2014). Structural basis of PcsB-mediated cell separation in *Streptococcus pneumoniae*. Nat. Commun. 5:3842.

6. Billini, M, Biboy, J, Kühn, J, Vollmer, W, Thanbichler, M (2019). A specialized MreB-dependent cell wall biosynthetic complex mediates the formation of stalk-specific peptidoglycan in *Caulobacter crescentus*. PLoS Genet. 15:e1007897.

7. Bochtler, M, Odintsov, SG, Marcyjaniak, M, Sabala, I (2004). Similar active sites in lysostaphins and D-Ala-D-Ala metallopeptidases. Protein Sci. 13:854–861.

8. Casadaban, MJ, Cohen, SN (1980). Analysis of gene control signals by DNA fusion and cloning in Escherichia coli. J. Mol. Biol. 138:179–207.

9. Cho, H, Uehara, T, Bernhardt, TG (2014). Beta-lactam antibiotics induce a lethal malfunctioning of the bacterial cell wall synthesis machinery. Cell 159:1300–1311.

10. Chodisetti, PK, Reddy, M (2019). Peptidoglycan hydrolase of an unusual cross-link cleavage specificity contributes to bacterial cell wall synthesis. Proc. Natl. Acad. Sci. USA 116:7825–7830.

11. Cook, J, Baverstock, TC, McAndrew, MBL, Stansfeld, PJ, Roper, DI, Crow, A (2020). Insights into bacterial cell division from a structure of EnvC bound to the FtsX periplasmic domain. Proc. Natl. Acad. Sci. USA 117:28355–28365.

12. Cserti, E, Rosskopf, S, Chang, Y-W, Eisheuer, S, Selter, L, Shi, J, Regh, C, Koert, U, Jensen, GJ, Thanbichler, M (2017). Dynamics of the peptidoglycan biosynthetic machinery in the stalked budding bacterium *Hyphomonas neptunium*. Mol. Microbiol. 103:875–895.

13. Curtis, PD, Brun, YV (2010). Getting in the loop. Microbiol. Mol. Biol. Rev. 74:13–41.

14. Dik, DA, Marous, DR, Fisher, JF, Mobashery, S (2017). Lytic transglycosylases: concinnity in concision of the bacterial cell wall. Crit. Rev. Biochem. Mol. Biol. 52:503–542.

15. Domínguez-Gil, T, Lee, M, Acebrón-Avalos, I, Mahasenan, KV, Hesek, D, Dik, DA, Byun, B, Lastochkin, E, Fisher, JF, Mobashery, S, et al. (2016). Activation by allostery in cell-wall remodeling by a modular membrane-bound lytic transglycosylase from Pseudomonas aeruginosa. Structure (London, England: 1993) 24: 1729-1741.

16. Dörr, T, Cava, F, Lam, H, Davis, BM, Waldor, MK (2013). Substrate specificity of an elongation-specific peptidoglycan endopeptidase and its implications for cell wall architecture and growth of *Vibrio cholerae*. Mol. Microbiol. 89:949–962.

17. Du, S, Lutkenhaus, J (2017). Assembly and activation of the *Escherichia coli* divisome. Mol. Microbiol. 105:177–187.

18. Dye, NA, Pincus, Z, Theriot, JA, Shapiro, L, Gitai, Z (2005). Two independent spiral structures control cell shape in *Caulobacter*. Proc. Natl. Acad. Sci. USA 102:18608–18613.

19. Ely, B (1991). Genetics of *Caulobacter crescentus*. Methods Enzymol. 204:372–384.

20. Emsley, P, Lohkamp, B, Scott, WG, Cowtan, K (2010). Features and development of Coot. Acta Crystallogr. D Biol. Crystallogr. 66:486–501.

21. Evans, PR, Murshudov, GN (2013). How good are my data and what is the resolution? Acta Crystallogr. D Biol. Crystallogr. 69:1204–1214.

22. Evans, R, O’Neill, M, Pritzel, A, Antropova, N, Senior, A, Green, T, Žídek, A, Bates, R, Blackwell, S, Yim, J, et al. (2022). Protein complex prediction with AlphaFold-Multimer. bioRxiv, DOI: 2021.2010.2004.463034.

23. Evinger, M, Agabian, N (1977). Envelope-associated nucleoid from *Caulobacter crescentus* stalked and swarmer cells. J. Bacteriol. 132:294–301.

24. Firczuk, M, Bochtler, M (2007). Folds and activities of peptidoglycan amidases. FEMS Microbiol. Rev. 31:676–691.

25. Gerding, MA, Liu, B, Bendezú, FO, Hale, CA, Bernhardt, TG, de Boer, PAJ (2009). Self-enhanced accumulation of FtsN at division sites and roles for other proteins with a SPOR domain (DamX, DedD, and RlpA) in Escherichia coli cell constriction. J. Bacteriol. 191:7383–7401.

26. Gitai, Z, Dye, N, Shapiro, L (2004). An actin-like gene can determine cell polarity in bacteria. Proc. Natl. Acad. Sci. USA 101:8643–8648.

27. Glauner, B (1988). Separation and quantification of muropeptides with high-performance liquid chromatography. Anal. Biochem. 172:451–464.

28. Goddard, TD, Huang, CC, Meng, EC, Pettersen, EF, Couch, GS, Morris, JH, Ferrin, TE (2018). UCSF ChimeraX: Meeting modern challenges in visualization and analysis. Protein Sci. 27:14–25.

29. Goley, ED, Comolli, LR, Fero, KE, Downing, KH, Shapiro, L (2010). DipM links peptidoglycan remodelling to outer membrane organization in Caulobacter. Mol. Microbiol. 77:56–73.

30. Goley, ED, Yeh, Y-C, Hong, S-H, Fero, MJ, Abeliuk, E, McAdams, HH, Shapiro, L (2011). Assembly of the Caulobacter cell division machine. Mol. Microbiol. 80:1680–1698.

31. Gurnani Serrano, CK, Winkle, M, Martorana, AM, Biboy, J, Morè, N, Moynihan, P, Banzhaf, M, Vollmer, W, Polissi, A (2021). ActS activates peptidoglycan amidases during outer membrane stress in *Escherichia coli*. Mol. Microbiol. 116:329–342.

32. Hartmann, R, van Teeseling, MCF, Thanbichler, M, Drescher, K (2018). BacStalk: a comprehensive and interactive image analysis software tool for bacterial cell biology. Mol. Microbiol. 114, 140–150.

33. Hashimoto, M, Ooiwa, S, Sekiguchi, J (2012). Synthetic lethality of the *lytE cwlO* genotype in *Bacillus subtilis* is caused by lack of D,L-endopeptidase activity at the lateral cell wall. J. Bacteriol. 194:796–803.

34. Heidrich, C, Templin, MF, Ursinus, A, Merdanovic, M, Berger, J, Schwarz, H, Pedro, MA, Höltje, JV (2001). Involvement of N-acetylmuramyl-L-alanine amidases in cell separation and antibiotic-induced autolysis of *Escherichia coli*. Mol. Microbiol. 41:167–178.

35. Heidrich, C, Ursinus, A, Berger, J, Schwarz, H, Holtje, J-V (2002). Effects of multiple deletions of murein hydrolases on viability, septum cleavage, and sensitivity to large toxic molecules in *Escherichia coli*. J. Bacteriol. 184:6093–6099.

36. Henrici, AT, Johnson, DE (1935). Studies of freshwater bacteria. J.Bacteriol. 30:61–93.

37. Höltje, JV (1998). Growth of the stress-bearing and shape-maintaining murein sacculus of *Escherichia coli*. Microbiol. Mol. Biol. Rev. 62:181–203.

38. Jacobs, C, Huang, LJ, Bartowsky, E, Normark, S, Park, JT (1994). Bacterial cell wall recycling provides cytosolic muropeptides as effectors for beta-lactamase induction. EMBO J. 13:4684–4694.

39. Jaqaman, K, Loerke, D, Mettlen, M, Kuwata, H, Grinstein, S, Schmid, SL, Danuser, G (2008). Robust single-particle tracking in live-cell time-lapse sequences. Nat. Methods 5:695–702.

40. Jung, A, Eisheuer, S, Cserti, E, Leicht, O, Strobel, W, Möll, A, Schlimpert, S, Kühn, J, Thanbichler, M (2015). Molecular toolbox for genetic manipulation of the stalked budding bacterium *Hyphomonas neptunium*. Appl. Environ. Microbiol. 81:736–744.

41. Kabsch, W (2010). Integration, scaling, space-group assignment and post-refinement. Acta Crystallogr. D Biol. Crystallogr. 66:133–144.

42. Kühn, J, Briegel, A, Mörschel, E, Kahnt, J, Leser, K, Wick, S, Jensen, GJ, Thanbichler, M (2010). Bactofilins, a ubiquitous class of cytoskeletal proteins mediating polar localization of a cell wall synthase in Caulobacter crescentus. EMBO J. 29:327–339.

43. Liebschner, D, Afonine, PV, Baker, ML, Bunkóczi, G, Chen, VB, Croll, TI, Hintze, B, Hung, LW, Jain, S, McCoy, AJ, et al. (2019). Macromolecular structure determination using X-rays, neutrons and electrons: recent developments in Phenix. Acta Crystallogr. D Struct. Biol. 75:861–877.

44. Lim, HC, Sher, JW, Rodriguez-Rivera, FP, Fumeaux, C, Bertozzi, CR, Bernhardt, TG (2019). Identification of new components of the RipC-FtsEX cell separation pathway of Corynebacterineae. PLoS Genet. 15:e1008284.

45. Lord, SJ, Velle, KB, Mullins, RD, Fritz-Laylin, LK (2020). SuperPlots: Communicating reproducibility and variability in cell biology. J. Cell Biol. 219:e202001064.

46. Lutkenhaus, J (2009). FtsN–trigger for septation. J.Bacteriol. 191:7381–7382.

47. Marblestone, J.G., Edavettal, S.C., Lim, Y., Lim, P., Zuo, X., and Butt, T.R. (2006). Comparison of SUMO fusion technology with traditional gene fusion systems: enhanced expression and solubility with SUMO. Prot. Sci. 15:182–189.

48. Martinelli, DJ, Pavelka, MS (2016). The RipA and RipB peptidoglycan endopeptidases are individually nonessential to *Mycobacterium smegmatis*. J. Bacteriol. 198:1464–1475.

49. Mavrici, D, Marakalala, MJ, Holton, JM, Prigozhin, DM, Gee, CL, Zhang, YJ, Rubin, EJ, Alber, T (2014). Mycobacterium tuberculosis FtsX extracellular domain activates the peptidoglycan hydrolase, RipC. Proc. Natl. Acad. Sci. USA 111:8037–8042.

50. Meier, EL, Daitch, AK, Yao, Q, Bhargava, A, Jensen, GJ, Goley, ED (2017). FtsEX-mediated regulation of the final stages of cell division reveals morphogenetic plasticity in *Caulobacter crescentus*. PLoS Genet. 13:e1006999.

51. Meiresonne, CE, Mertens, LMY, Chertkova, AO, Goedhart, J, den Blaauwen, T (2019). Superfolder mTurquoise2^ox^ optimized for the bacterial periplasm allows high efficiency in vivo FRET of cell division antibiotic targets. Mol. Microbiol. 111: 1025–1038.

52. Meisenzahl, AC, Shapiro, L, Jenal, U (1997). Isolation and characterization of a xylose-dependent promoter from *Caulobacter crescentus*. J. Bacteriol. 179:592–600.

53. Meisner, J, Montero Llopis, P, Sham, L-T, Garner, E, Bernhardt, TG, Rudner, DZ (2013). FtsEX is required for CwlO peptidoglycan hydrolase activity during cell wall elongation in *Bacillus subtilis*. Mol. Microbiol. 89:1069–1083.

54. Mistry, J, Chuguransky, S, Williams, L, Qureshi, M, Salazar, GA, Sonnhammer, ELL, Tosatto, SCE, Paladin, L, Raj, S, Richardson, LJ, et al. (2021). Pfam: The protein families database in 2021. Nucleic Acids Res. 49:D412–D419.

55. Möll, A, Dörr, T, Alvarez, L, Chao, MC, Davis, BM, Cava, F, Waldor, MK (2014). Cell separation in *Vibrio cholerae* is mediated by a single amidase whose action is modulated by two nonredundant activators. J. Bacteriol. 196:3937–3948.

56. Möll, A, Schlimpert, S, Briegel, A, Jensen, GJ, Thanbichler, M (2010). DipM, a new factor required for peptidoglycan remodelling during cell division in *Caulobacter crescentus*. Mol. Microbiol. 77:90–107.

57. Möll, A, Thanbichler, M (2009). FtsN-like proteins are conserved components of the cell division machinery in proteobacteria. Mol. Microbiol. 72:1037–1053.

58. Mukherjee, A, Lutkenhaus, J (1994). Guanine nucleotide-dependent assembly of FtsZ into filaments. J. Bacteriol. 176:2754–2758.

59. Müller, P, Ewers, C, Bertsche, U, Anstett, M, Kallis, T, Breukink, E, Fraipont, C, Terrak, M, Nguyen-Distèche, M, Vollmer, W (2007). The essential cell division protein FtsN interacts with the murein (peptidoglycan) synthase PBP1B in *Escherichia coli*. J. Biol. Chem. 282:36394–36402.

60. Murshudov, GN, Skubák, P, Lebedev, AA, Pannu, NS, Steiner, RA, Nicholls, RA, Winn, MD, Long, F, Vagin, AA (2011). REFMAC5 for the refinement of macromolecular crystal structures. Acta Crystallogr. D Biol. Crystallogr.y 67:355–367.

61. Ortiz, C, Natale, P, Cueto, L, Vicente, M (2016). The keepers of the ring. FEMS Microbiol. Rev. 40:57–67.

62. Oviedo-Bocanegra, LM, Hinrichs, R, Rotter Daniel Andreas O, Dersch, S, Graumann, PL (2021). Single molecule/particle tracking analysis program SMTracker 2.0 reveals different dynamics of proteins within the RNA degradosome complex in *Bacillus subtilis*. Nucleic Acids Res. 49:e112.

63. Pazos, M, Peters, K, Vollmer, W (2017). Robust peptidoglycan growth by dynamic and variable multi-protein complexes. Curr. Opin. Microbiol. 36:55–61.

64. Peters, NT, Dinh, T, Bernhardt, TG (2011). A fail-safe mechanism in the septal ring assembly pathway generated by the sequential recruitment of cell separation amidases and their activators. J. Bacteriol. 193:4973–4983.

65. Peters, NT, Morlot, C, Yang, DC, Uehara, T, Vernet, T, Bernhardt, TG (2013). Structure-function analysis of the LytM domain of EnvC, an activator of cell wall remodelling at the *Escherichia coli* division site. Mol. Microbiol. 89:690–701.

66. Poggio, S, Takacs, CN, Vollmer, W, Jacobs-Wagner, C (2010). A protein critical for cell constriction in the Gram-negative bacterium *Caulobacter crescentus* localizes at the division site through its peptidoglycan-binding LysM domains. Mol. Microbiol. 77:74–89.

67. Poindexter, JS (1981). The caulobacters. Microbiol. Rev. 45:123–179.

68. Poindexter, JS (1964). Biological properties and classification of the *Caulobacter* group. Bacteriol. Rev. 28:231–295.

69. Postma, M, Goedhart, J (2019). PlotsOfData—A web app for visualizing data together with their summaries. PLOS Biol. 17:e3000202.

70. Reinscheid, DJ, Gottschalk, B, Schubert, A, Eikmanns, BJ, Chhatwal, GS (2001). Identification and molecular analysis of PcsB, a protein required for cell wall separation of group B streptococcus. J. Bacteriol. 183:1175–1183.

71. Rued, BE, Alcorlo, M, Edmonds, KA, Martínez-Caballero, S, Straume, D, Fu, Y, Bruce, KE, Wu, H, Håvarstein, LS, Hermoso, JA, et al. (2019). Structure of the large extracellular loop of FtsX and its interaction with the essential peptidoglycan hydrolase PcsB in *Streptococcus pneumoniae*. mBio 10:e02622.

72. Schindelin, J, Arganda-Carreras, I, Frise, E, Kaynig, V, Longair, M, Pietzsch, T, Preibisch, S, Rueden, C, Saalfeld, S, Schmid, B, et al. (2012). Fiji: an open-source platform for biological-image analysis. Nat. Methods 9:676-682.

73. Schleifer, KH, Kandler, O (1972). Peptidoglycan types of bacterial cell walls and their taxonomic implications. Bacteriol Rev. 36:407–477.

74. Schmidt, KL, Peterson, ND, Kustusch, RJ, Wissel, MC, Graham, B, Phillips, GJ, Weiss, DS (2004). A predicted ABC transporter, FtsEX, is needed for cell division in *Escherichia coli*. J. Bacteriol. 186:785–793.

75. Sham, L-T, Barendt, SM, Kopecky, KE, Winkler, ME (2011). Essential PcsB putative peptidoglycan hydrolase interacts with the essential FtsXSpn cell division protein in *Streptococcus pneumoniae* D39. Proc. Natl. Acad. Sci. USA 108:E1061–1069.

76. Singh, SK, SaiSree, L, Amrutha, RN, Reddy, M (2012). Three redundant murein endopeptidases catalyse an essential cleavage step in peptidoglycan synthesis of *Escherichia coli* K12. Mol. Microbiol. 86:1036–1051.

77. Strobel, W, Möll, A, Kiekebusch, D, Klein, KE, Thanbichler, M (2014). Function and localization dynamics of bifunctional penicillin-binding proteins in *Caulobacter crescentus*. J. Bacteriol. 196:1627–1639.

78. Thanbichler, M, Iniesta, AA, Shapiro, L (2007). A comprehensive set of plasmids for vanillate- and xylose-inducible gene expression in *Caulobacter crescentus*. Nucleic Acids Res. 35:e137.

79. Thanbichler, M, Shapiro, L (2006). MipZ, a spatial regulator coordinating chromosome segregation with cell division in *Caulobacter*. Cell 126:147–162.

80. Tsai, JW, Alley, MR (2001). Proteolysis of the *Caulobacter* McpA chemoreceptor is cell cycle regulated by a ClpX-dependent pathway. J. Bacteriol. 183:5001–5007.

81. Tsang, M-J, Yakhnina, AA, Bernhardt, TG (2017). NlpD links cell wall remodeling and outer membrane invagination during cytokinesis in *Escherichia coli*. PLoS Genet. 13:e1006888.

82. Tyanova, S, Temu, T, Sinitcyn, P, Carlson, A, Hein, MY, Geiger, T, Mann, M, Cox, J (2016). The Perseus computational platform for comprehensive analysis of (prote)omics data. Nat. Methods 13:731–740.

83. Typas, A, Banzhaf, M, Gross, CA, Vollmer, W (2011). From the regulation of peptidoglycan synthesis to bacterial growth and morphology. Nat. Rev. Microbiol. 10:123–136.

84. Uehara, T, Dinh, T, Bernhardt, TG (2009). LytM-domain factors are required for daughter cell separation and rapid ampicillin-induced lysis in *Escherichia coli*. J. Bacteriol. 191:5094–5107.

85. Uehara, T, Parzych, KR, Dinh, T, Bernhardt, TG (2010). Daughter cell separation is controlled by cytokinetic ring-activated cell wall hydrolysis. EMBO J. 29:1412–1422.

86. Vollmer, W, Blanot, D, Pedro, MA (2008). Peptidoglycan structure and architecture. FEMS Microbiol. Rev. 32:149–167.

87. Wagner, JK, Galvani, CD, Brun, YV (2005). *Caulobacter crescentus* requires RodA and MreB for stalk synthesis and prevention of ectopic pole formation. J. Bacteriol. 187:544–553.

88. Weidel, W, Frank, H, Martin, HH (1960). The rigid layer of the cell wall of *Escherichia coli* strain B. J. Gen. Microbiol. 22:158–166.

89. Weiss, DS (2015). Last but not least: new insights into how FtsN triggers constriction during *Escherichia coli* cell division. Mol. Microbiol. 95:903–909.

90. Yahashiri, A, Jorgenson, MA, Weiss, DS (2015). Bacterial SPOR domains are recruited to septal peptidoglycan by binding to glycan strands that lack stem peptides. Proc. Natl. Acad. Sci. USA 112:11347–11352.

91. Yahashiri, A, Jorgenson, MA, Weiss, DS (2017). The SPOR domain, a widely conserved peptidoglycan binding domain that targets proteins to the site of cell division. J. Bacteriol. 199:e00118–17.

92. Yakhnina, AA, McManus, HR, Bernhardt, TG (2015). The cell wall amidase AmiB is essential for *Pseudomonas aeruginosa* cell division, drug resistance and viability. Mol. Microbiol. 97:957–973.

93. Yang, DC, Tan, K, Joachimiak, A, Bernhardt, TG (2012). A conformational switch controls cell wall-remodelling enzymes required for bacterial cell division. Mol. Microbiol. 85:768–781.

94. Yang, L-C, Gan, Y-L, Yang, L-Y, Jiang, B-L, Tang, J-L (2018). Peptidoglycan hydrolysis mediated by the amidase AmiC and its LytM activator NlpD is critical for cell separation and virulence in the phytopathogen *Xanthomonas campestris*. Mol. Plant Pathol. 19:1705–1718.

95. Zielińska, A, Billini, M, Möll, A, Kremer, K, Briegel, A, Izquierdo Martinez, A, Jensen, GJ, Thanbichler, M (2017). LytM factors affect the recruitment of autolysins to the cell division site in *Caulobacter crescentus*. Mol. Microbiol. 106:419–438.

## Supplementary references

96. An, DR, Im, HN, Jang, JY, Kim, HS, Kim, J, Yoon, HJ, Hesek, D, Lee, M, Mobashery, S, Kim, S-J, et al. (2016). Structural basis of the heterodimer formation between cell shape-determining proteins Csd1 and Csd2 from *Helicobacter pylori*. PLoS One 11:e0164243.

97. Billini, M, Biboy, J, Kühn, J, Vollmer, W, Thanbichler, M (2019). A specialized MreB-dependent cell wall biosynthetic complex mediates the formation of stalk-specific peptidoglycan in *Caulobacter crescentus*. PLoS Genet 15:e1007897.

98. Cook, J, Baverstock, TC, McAndrew, MBL, Stansfeld, PJ, Roper, DI, Crow, A (2020). Insights into bacterial cell division from a structure of EnvC bound to the FtsX periplasmic domain. Proc Natl Acad Sci USA 117:28355–28365.

99. Evans, R, O’Neill, M, Pritzel, A, Antropova, N, Senior, A, Green, T, Žídek, A, Bates, R, Blackwell, S, Yim, J, et al. (2022). Protein complex prediction with AlphaFold-Multimer. bioRxiv, DOI: 10.1101/2021.10.04.463034.

100. Evinger, M, Agabian, N (1977). Envelope-associated nucleoid from *Caulobacter crescentus* stalked and swarmer cells. J Bacteriol 132:294–301.

101. Grabowska, M, Jagielska, E, Czapinska, H, Bochtler, M, Sabala, I (2015). High resolution structure of an M23 peptidase with a substrate analogue. Sci Rep 5:14833.

102. Meisner, J, Maehigashi, T, André, I, Dunham, CM, Moran, CP (2012). Structure of the basal components of a bacterial transporter. Proc Natl Acad Sci USA 109:5446–5451.

103. Mistry, J, Chuguransky, S, Williams, L, Qureshi, M, Salazar, GA, Sonnhammer, ELL, Tosatto, SCE, Paladin, L, Raj, S, Richardson, LJ, et al. (2021). Pfam: The protein families database in 2021. Nucleic Acids Res 49:D412–D419.

104. Möll, A, Schlimpert, S, Briegel, A, Jensen, GJ, Thanbichler, M (2010). DipM, a new factor required for peptidoglycan remodelling during cell division in *Caulobacter crescentus*. Mol Microbiol 77:90–107.

105. Möll, A, Thanbichler, M (2009). FtsN-like proteins are conserved components of the cell division machinery in proteobacteria. Mol Microbiol 72:1037–1053.

106. Shin, J-H, Sulpizio, AG, Kelley, A, Alvarez, L, Murphy, SG, Fan, L, Cava, F, Mao, Y, Saper, MA, Dörr, T (2020). Structural basis of peptidoglycan endopeptidase regulation. Proc Natl Acad Sci USA 117:11692–11702.

107. Thanbichler, M, Iniesta, AA, Shapiro, L (2007). A comprehensive set of plasmids for vanillate- and xylose-inducible gene expression in *Caulobacter crescentus*. Nucleic Acids Res 35:e137.

108. Zielińska, A, Billini, M, Möll, A, Kremer, K, Briegel, A, Izquierdo Martinez, A, Jensen, GJ, Thanbichler, M (2017). LytM factors affect the recruitment of autolysins to the cell division site in *Caulobacter crescentus*. Mol Microbiol 106:419–438.

